# Single-Cell Transcriptomics Reveals the Molecular Logic Underlying Ca^2+^ Signaling Diversity in Human and Mouse Brain

**DOI:** 10.1101/2024.04.26.591400

**Authors:** Ibrahim Al Rayyes, Lauri Louhivuori, Ivar Dehnisch Ellström, Erik Smedler, Per Uhlén

## Abstract

The calcium ion (Ca^2+^) is a ubiquitous intracellular signaling molecule that plays a critical role in the adult and developing brain. However, the principles governing the specificity of Ca^2+^ signaling remain unresolved. In this work, we comprehensively analyzed the Ca^2+^ signaling transcriptome in the adult mouse brain and developing human brain. We found that neurons form non-stochastic Ca^2+^-states that are reflective of their cell types and functionality, with evidence suggesting that the diversity is driven by lineage-specific developmental changes. Focusing on the neocortical development, we reveal that an unprecedented number of Ca^2+^ genes are tightly regulated and evolutionarily conserved, capturing functionally driven differences within radial glia and neuronal progenitors. In summary, our study provides an in-depth understanding of the cellular and temporal diversity of Ca^2+^ signaling and suggests that Ca^2+^ signaling is dynamically tailored to specific cell states.

**One Sentence Summary:** The expression of Ca^2+^ signaling genes is finely tuned to cellular states, reflecting a spectrum of differences that range from lineage specificity to subtle functional distinctions within cortical radial glia.

## Main Text

Calcium ions (Ca^2+^) are versatile intracellular signaling messengers that regulate most neuronal functions, including synaptic plasticity, neurotransmitter release, excitability, and activity-dependent gene transcription (*1–3*). Through interactions with developmentally important signaling pathways, such as Notch- and Wnt-signaling (*4, 5*), morphogens (*6*), and transcription factors (TFs) (*7*), Ca^2+^ signaling is known to regulate nearly every aspect of neurogenesis, synaptogenesis and experience-dependent plasticity across multiple regions, including the neocortex, cerebellum, dentate gyrus, and olfactory bulb (*8, 9*). This versatility is enabled by the Ca^2+^ signaling toolkit, a large repertoire of proteins that control, transduce, and act on Ca^2+^ signals to generate downstream effects. Given the diversity in neuronal functions, firing properties, and connectivity, as well as the complex spatiotemporal dynamics of morphogens and patterning genes that ultimately guide developmental programs across the brain, the involvement of Ca^2+^ signaling in these processes raises the question of how different outputs from Ca^2+^ signals are achieved.

One hypothesis suggests that the tissue-specific expression of subsets of genes that encode the Ca^2+^ signaling toolkit, here referred to as the Ca^2+^ signaling transcriptome, could enable cells to achieve and maintain functional specificity (*1, 10*). Whether this hypothesis applies to the brain remains an unexplored question. Nevertheless, there are examples, such as the utilization of *PVALB* and *CALB2* to identify specific sub-populations of midbrain dopaminergic and cortical GABAergic neurons (*11, 12*), as well as differences between the cerebellum and dentate gyrus in regards to how Ca^2+^ mediated mechanisms drive long-term potentiation and depression (*13*) that may reflect a general pattern of transcriptional cell- and region-specificity in Ca^2+^ signaling. Understanding the molecular principles underlying the specificity of Ca^2+^ signaling could have significant implications for our comprehension of Ca^2+^-driven processes, including the regulation of neurogenesis across different brain regions.

Until recently, technological limitations have constrained comprehensive studies of the Ca^2+^ signaling transcriptome, resulting in poor understanding of the diversity of transcriptional Ca^2+^-states. With the rapid emergence of single-cell RNA sequencing (scRNA-seq), an unprecedented capacity for studying the temporal and cellular heterogeneity of Ca^2+^ signaling in the human and mouse brain now exists.

## Results

### scRNA-seq reveals an architecture of distinct Ca^2+^-states across the nervous system

To gain a comprehensive understanding of the Ca^2+^ signaling transcriptome in the nervous system, we curated a list of Ca^2+^ genes (*14*) and analyzed their expressions in the adult mouse nervous system by Zeisel et al. (*15*). We used single-cell Hierarchical Poisson Factorization (scHPF) to decompose the counts into 27 discrete Ca^2+^ signaling modules (Fig. 1A, Fig. S1A-C), with a module score attributed to every cell and gene. We next devised a clustering strategy that utilized the cell-module scores and identified 28 unique cellular Ca^2+^ states that largely reflected the underlying cell type architecture (Fig. 1B; Table S1), with each Ca^2+^ state enriched for at least one Ca^2+^ signaling module (Fig. S1E; Fig. S2). To evaluate the robustness and potential technical batch effects, as well as to determine whether the Ca^2+^ states represented biologically meaningful clusters or mere artifacts of general gene expression differences across cell types, we employed a two-pronged approach. First, we ran multiple iterations using an equivalent number of random genes. Then, we trained random forest classifiers to predict both Ca^2+^-states and cell types. We found that the random genes did not recapitulate the Ca^2+^-driven clustering and that the Ca^2+^-based classifiers were superior in predicting Ca^2+^-states and cell types (Fig. S3), indicating that Ca^2+^ genes are biologically informative.

**Fig. 1.**
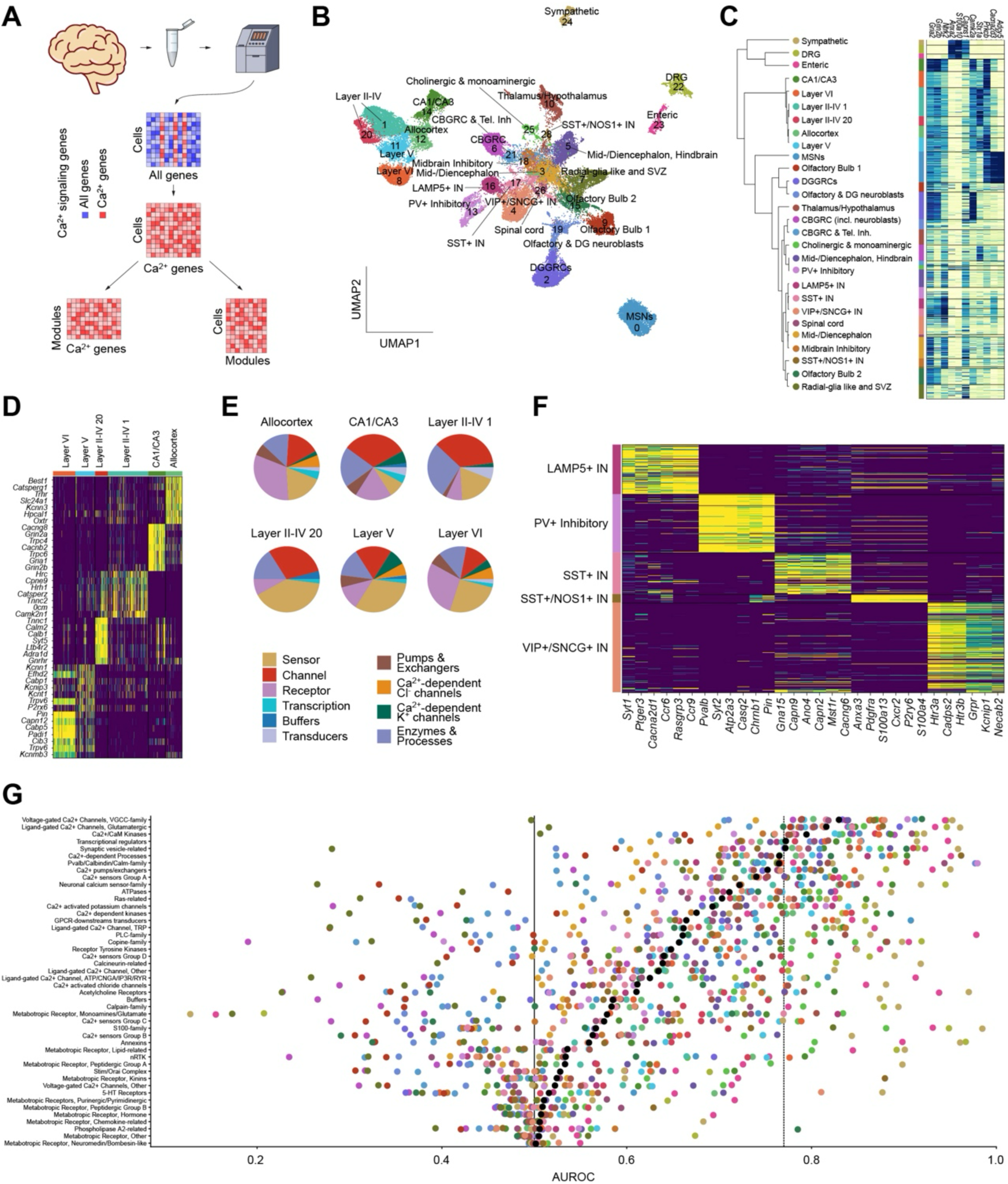
Characterization of single-cell Ca^2+^-states across the mouse nervous system. (**A**) Schematic overview of the analysis workflow. All non-Ca^2+^ genes were removed, and the remaining unique molecular identifier (UMI) counts were decomposed using scHPF. The gene and cell scores were used for downstream clustering and gene expression analysis. (**B**) A Ca^2+^-based UMAP embedding of the 63,507 cells from Zeisel et al. (*15*), clustered into 27 Ca^2+^-states. (**C**) *Left:* Dendrogram constructed from the high-dimensional UMAP plot illustrating the similarities between Ca^2+^-states. *Right:* Heatmap showing examples of genes related to the successive splits across the mouse nervous system. Rows show single cells, grouped by Ca^2+^-states. (**D**) Heatmap of the telencephalic excitatory neuronal Ca^2+^-states, illustrating the differential enrichments of Ca^2+^ genes. (**E**) Pie charts of the gene classes of the top 50 genes per telencephalic excitatory neuronal Ca^2+^-state. (**F**) Same as (**D**), but for telencephalic interneurons. (**G**) Scatter plots of AUROC scores for all 46 functional groups, ordered by mean. Colors indicate Ca^2+^-states.

To link the Ca^2+^ states directly to genes, we calculated cell-gene scores from the cell-module and gene-module scores (Table S2; Fig. S4A; Methods). A dendrogram constructed on a high-dimensional uniform manifold approximation and projection (UMAP) plot showed that the first split in Ca^2+^ states was between the central nervous system (CNS) and peripheral nervous system (PNS) neurons, marked by differences in genes such as *Gria2* and *Grin2b* or *Anxa2* and *S100a10* (Fig. 1C). This was followed by a split between the telencephalic excitatory neurons and medium spiny neurons (MSN), enriched for *Camk2a, Stx1a* and *Cacna2d3, Adcy5,* respectively. The telencephalic excitatory neurons formed six Ca^2+^ states with divisions into neocortical layers II-IV, V, and VI, allocortex, and hippocampal CA1/CA3 neurons. Each Ca^2+^ state exhibited unique patterns on a gene (Fig. 1D) and gene family level (Fig. 1E), with particularly clear differences in the proportion of Receptors (e.g., *Adra1d, Oxtr*), Sensors (e.g., *Calb1, Calm2*), Channels (e.g., *Grin2b, Gria1*), and Enzymes & Processes (e.g., *Camk2n1*, *Capn9*).

Moreover, we found that the telencephalic interneurons formed distinctive Ca^2+^ states that largely adhered to the major interneuron cell types (Fig. S4B), with one Ca^2+^ state shared by the VIP+ and SNCG+ interneurons (Fig. S4C). Notably, the separation was driven by differences in a range of Ca^2+^ genes including *Cacna2d1*, *Rasgrp3, Atp2a3, Cacng6*, *Anxa3*, *S100a13*, *Htr3a*, and *Htr3b,* illustrating that the differential expression of Ca^2+^ genes in interneurons extends beyond *Pvalb* and *Calb2,* which have commonly been used to classify interneurons (Fig. 1F). Notably, the PV+ Inhibitory Ca^2+^ state also included non-telencephalic inhibitory neurons, such as the cerebellar molecular layer interneurons. Additionally, one of the two SST+ Ca^2+^ states was shared with the dopaminergic olfactory interneurons, suggesting that interneurons of different developmental origins overlap in their utilization of Ca^2+^ genes. Our findings show that the different classes of interneurons express distinct subsets of the Ca^2+^ signaling transcriptome, thereby suggesting a generalizable specification of how Ca^2+^ signaling is used by interneurons.

We then categorized the Ca^2+^ genes based on their function (Table S3) and evaluated how each functional group contributed to transcriptomic heterogeneity. The “Voltage-gated Ca^2+^ channels, VGCC-family”, “Ligand-gated Ca^2+^ channels, Glutamatergic”, Ca^2+^/CaM kinases”, “Transcriptional regulators”, and “Synaptic vesicle-related” contributed highly across the entire dataset, indicating that these sets are differentially enriched across the Ca^2+^ states (Fig. 1G). Numerous functional groups, including the “S100-family” and “Annexins”, performed strongly in predicting the DRG, Sympathetic, ENS Ca^2+^ states despite a lower overall performance (Fig. S4D), suggesting that they were less discernible within the CNS, and that the largest axis of variation in the Ca^2+^ signaling transcriptome was between the CNS and PNS. Together, our results illustrate, for the first time, that Ca^2+^ genes are heterogeneously expressed across the brain in a function- and cell type-dependent manner, raising the possibility that neurons are transcriptionally primed towards specific cell responses or functions.

### The cellular Ca^2+^ states capture activity-driven developmental changes within cortical cell types

Given the many functional roles of Ca^2+^ signaling, we next asked whether the Ca^2+^ states may convey information about the functionality of cells unrelated to their taxonomic identities. To address this question, we focused on *Ca^2+^ state 1* and *20,* given that they were dominated by the same cell types: *TEGLU7, TEGLU8*, and *TEGLU12* (Fig. 2A). First, we confirmed that the cells were sourced from multiple donors sequenced across several experiments (Fig. S5A), thus making it less likely that the differences in Ca^2+^ module enrichment (Fig. 2B) were caused by batch effects. Next, we confirmed that there were no apparent differences in the expressions of cell type markers between *Ca^2+^ state 1* and *20* (Fig. S5B).

**Fig. 2.**
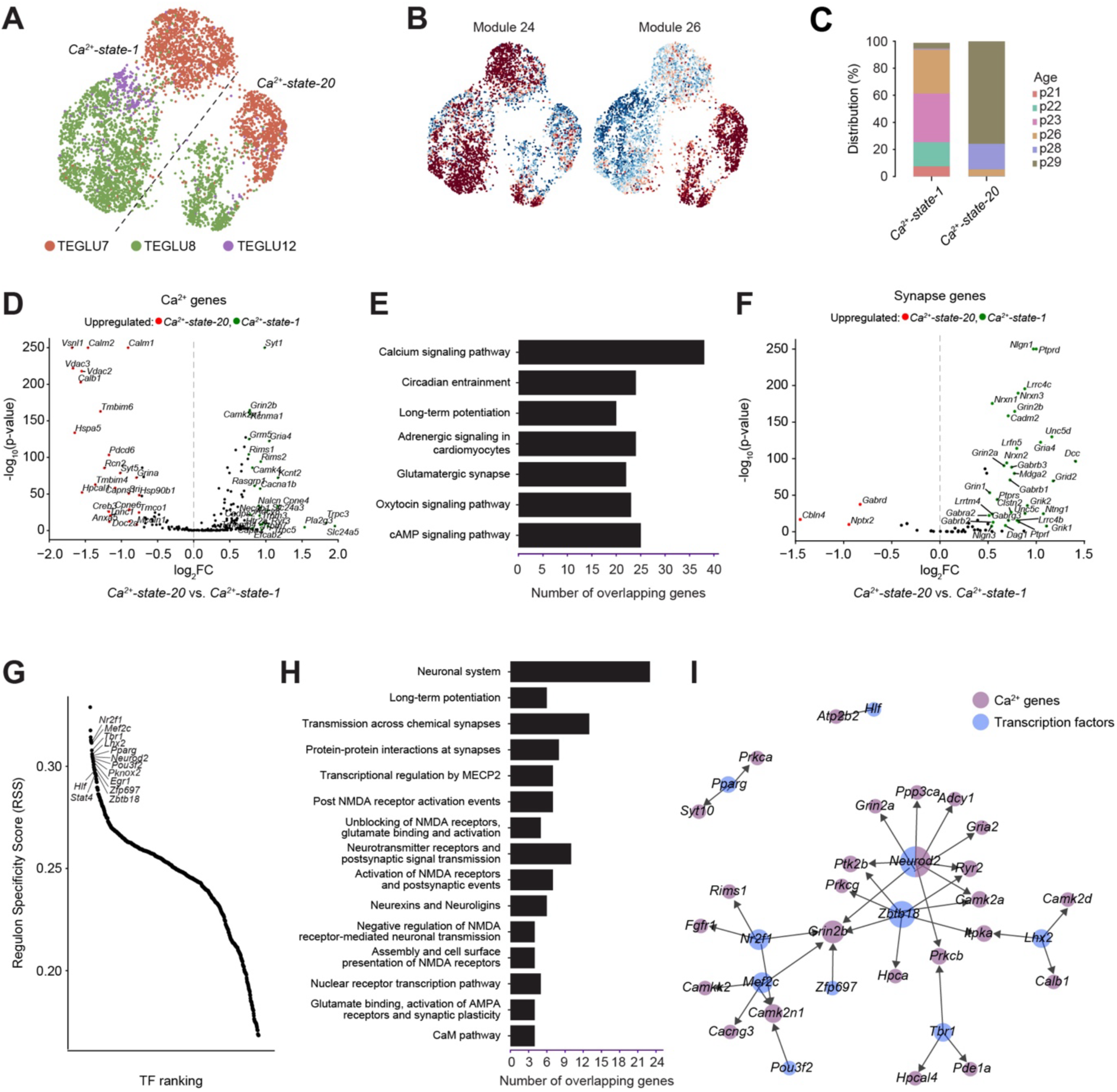
Ca^2+^-states capture functional activity-driven heterogeneity within identical neocortical cell types. (**A**) UMAP embedding illustrating cell types found in *Ca^2+^-state-1* and *Ca^2+^-state-20*, indicated by colors. Dashed lines illustrate the borders separating the two Ca^2+^-states. (**B**) Feature plots illustrating the differential enrichments of *module 24* and *module 26*. (**C**) Stacked bar plots showing the age distributions within the two Ca^2+^-states. (**D**) Volcano plot of Ca^2+^ genes, illustrating the differential expressions across the two Ca^2+^-states. (**E**) KEGG pathway enrichment in *Ca^2+^-state-1* compared to *Ca^2+^-state-20*, ranked by *p*-values. (**F**) Same as (**D**), but for genes related to synapse remodeling, adapted from Südhof (*17*). (**G**) Top unique TFs in *Ca^2+^-state-1* ranked by the regulon specificity score (RSS), with respect to the entire mouse dataset. (**H**) Reactome pathway analysis of enriched regulons in *Ca^2+^-state-1* compared to *Ca^2+^-state-20.* (**I**) Network illustrating TFs that are predicted to directly upregulate Ca^2+^ genes.

To further understand the discrepancy between the Ca^2+^ states and the originally defined taxonomy, we analyzed the relationship between Ca^2+^ states and the global transcriptome. Notably, a linear discriminant analysis of the principal components (PCs) of the global transcriptome revealed that the cells adhered to their Ca^2+^ states based on a few PCs. As the number of PCs increased, a transition towards their taxonomic cell types was observed (Fig. S5C-D), thereby masking the Ca^2+^ states and illustrating that the Ca^2+^ states reflect subtle differences within cell types. In particular, we found that cells in *Ca^2+^ state 1* were primarily aged P21-P23, whereas those in *Ca^2+^ state 20* were predominantly P29 (Fig. 2C). Differential expression analysis of the Ca^2+^ genes showed that *Ca^2+^ state 1* exhibited an upregulation of the VGCC, NMDAR, and CAMKII-related genes, amongst others (Fig. 2D; Fig. S5E; Table S4). Furthermore, gene ontology analysis indicated an overall enrichment for Ca^2+^ genes belonging to “Long-term potentiation”, “Glutamatergic synapse”, and CaMP signaling pathway” (Fig. 2E). Given that the P21-P23 align with the “critical period” – a phase marked by heightened potential for experience-driven synaptic plasticity and maturation during postnatal development (*16*) – these findings suggest that the division into two Ca^2+^ states might reflect variations in activity-dependent synaptic plasticity. Consistent with this, differential expression analysis of genes known to drive synapse formation and remodeling, adapted from Südhof (*17*) (Table S5), revealed that many were upregulated in *Ca^2+^ state 1* (Fig. 2F), including Neurexins (*Nrxn1*, *Nrxn2, Nrxn3*), Neuroligins (*Nlgn1, Nlgn2*, *Nlgn3*), SynCAM2 (*Cadm2*), LAR-type receptor tyrosine phosphatases (*Ptprs*, *Ptprd, Ptprf*), and GluD2 (*Grid2*). Stratifying by cell type preserved the observed differences between *Ca^2+^ state 1* and *20* (Fig. S5F). Additionally, we found that these expression differences were consistently replicated across other neocortical cell types exhibiting similar age distributions (Fig. S5G).

Finally, to further validate our findings, we explored if the observed transcriptional differences between *Ca^2+^ state 1* and *20* were reflected in differences in TF activities. As expected, we found that the predicted TF activities of the two Ca^2+^ states were highly correlated (Spearman correlation = 0.87, Fig. S6A), reflecting their overall similarities. Next, we calculated specificity scores for the two Ca^2+^ states across the entire dataset. Of the top 20 TFs, seven were shared, including the upper-layer markers *Emx1* and *Cux1* (Fig. S6B. Furthermore, *Ca^2+^ state 1* was highly specific for *Mef2c, Nr2f1*, and *Zbtb18*, amongst others (Fig. 2G) which have previously been implicated in regulating activity-dependent synapse formation (*18–20*). Gene ontology analysis of the thirteen non-shared TFs and their inferred targets showed that the TFs overall converged on pathways and genes related to synaptic plasticity and remodeling (Fig. 2H; Fig. S6C-D), involving the upregulation of Ca^2+^ genes such as *Camk2n1*, *Camk2a*, and *Grin2b* (Fig. 2I). These results thus exemplify the potential of Ca^2+^ gene analysis to provide deeper biological insights, revealing subtle yet significant differences among highly similar cells.

### Postnatal remodeling of cellular Ca^2+^ states follow lineage-specific neurogenesis trajectories

Notably, the dentate gyrus (DG) and olfactory (OL) radial glial cell (RGC)-like cells formed a common Ca^2+^ state together with the sub-ventricular zone progenitors. Furthermore, the UMAP plots showed that this Ca^2+^ state bordered the DG and OL neuroblasts, which in turn were juxtaposed with the DG granule neurons and OL neurons (Fig. 1B), mimicking lineage trajectories. The increasing divergence with maturation warranted a closer analysis of the dynamics of the Ca^2+^ states during postnatal development. Accordingly, we first refined the separation of Ca^2+^ states by re-iterating the Ca^2+^- driven clustering on a subset of the dataset restricted to the neural progenitors, neuroblasts, and their mature counterparts: DG granule neurons, OL inhibitory neurons, and cerebellar (CB) granule neurons. This refined the separation of the progenitor Ca^2+^ state (Fig. S7A) and aligned the cells along three lineage trajectories (Fig. 3A) with differential enrichment of Ca^2+^ modules across the manifold (Fig. S7B). Next, we performed RNA velocity and cell fate analyses exclusively on the Ca^2+^ genes to contrast the dynamics of Ca^2+^ genes within and between trajectories. The three types of neurons appeared as terminal Ca^2+^ states, and the early RG-like cells were the only initial Ca^2+^ state (Fig. 3B; Fig. S7C). Each terminal Ca^2+^ state correlated with different lineage-determining genes, such as *Ncald* and *Ppp3ca* (DG), *Prkca* and *Cpne4* (OL), *Camk4* and *Cacna1a* (CB), whereas the RG-like Ca^2+^ state correlated with *Ntsr2, Sparc, Notch1*, *Ednrb* and several S100A-members (Fig. 3C, Fig. S7D-E). Using the DG trajectory as an example, we noted a gradual increase in the expression of terminal genes within the immature *DGNBL2* population that correlated with pseudotime (Fig. 3D), highlighting the potential existence of a gradient of functional maturity in developing neurons. Furthermore, ordering the lineage-related genes by their pseudotemporal kinetics revealed that the global transcriptional Ca^2+^ dynamics differed between the three trajectories (Fig. 3E). As expected from the UMAP plots, the DG and OL lineages had many overlapping genes compared to the CB (36 vs. 10 and 11, respectively), and no Ca^2+^ genes were shared across all three (Fig. 3F). Bifurcation analysis showed that a total of 28 genes drove the bifurcation between the DG and OL lineages, including members of the KCNIP-family of Ca^2+^-dependent transcription factors (*Kcnip1* and *Kcnip4*), *Camk2a, Camk2b,* and the VGCCs, that were shared between the two branches (Fig. 3G-H), suggesting that Ca^2+^ signaling genes are significantly influential in controlling the determination of cellular fate.

**Fig. 3.**
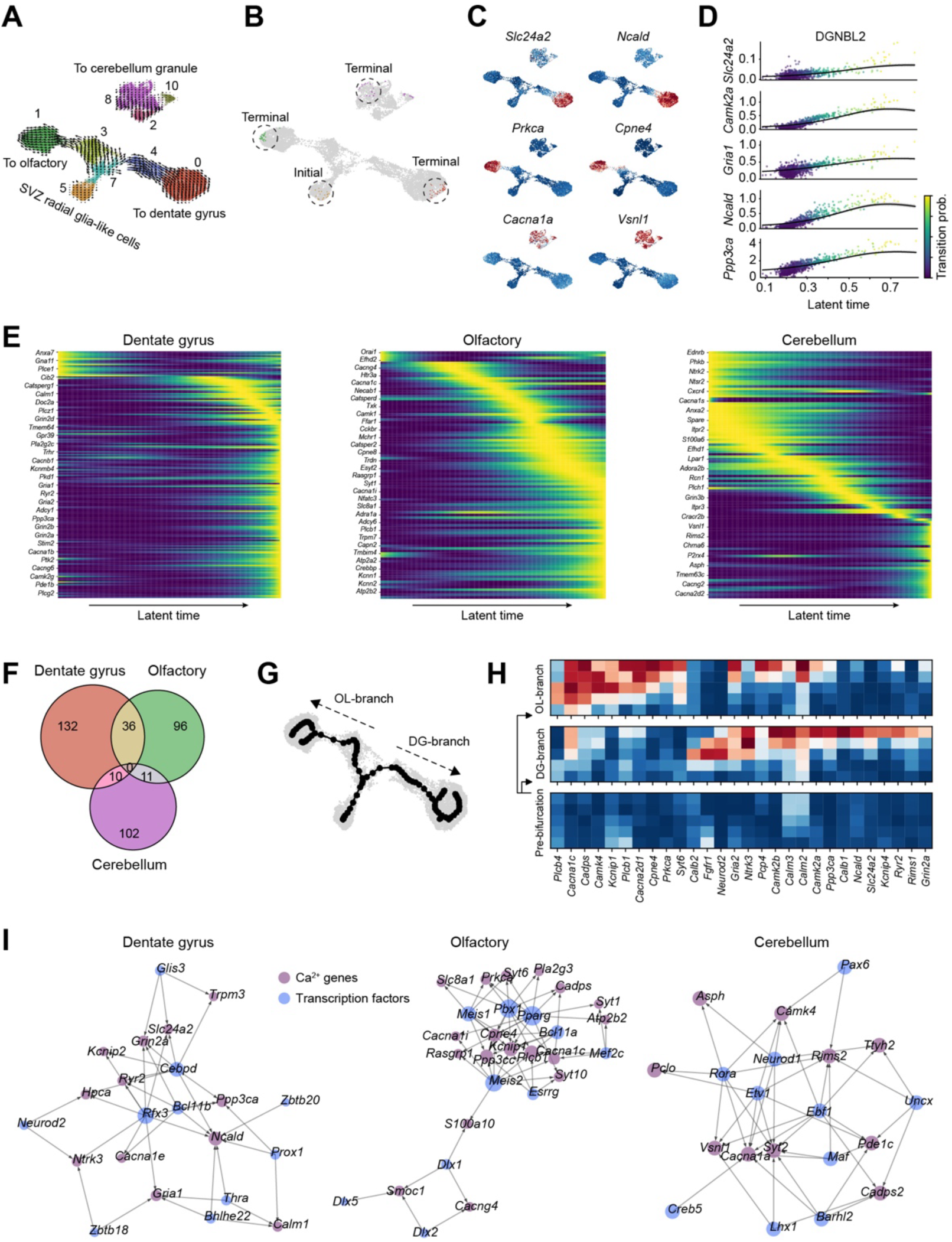
Postnatal remodeling of Ca^2+^-states follow lineage-specific neurogenesis trajectories. (**A**) UMAP plot illustrating the divergent trajectories based on transcriptomic changes in Ca^2+^ signaling genes. Arrows indicate Ca^2+^-based velocity vectors. (**B**) Lineage-inference analysis using CellRank on Ca^2+^ genes indicate that the immature RG-like cells constitute the earliest state, and that three terminal-states exist: the DGGRCs, OL, and CBGRCs. (**C**) Feature plots showing the expression of the most highly correlating Ca^2+^ genes with each of the three lineages. (**D**) Scatterplot showing the gradient of increased expression of end-stage dentate gyrus Ca^2+^ genes within the DGNBL2 population. Colors indicate absorption probability. (**E**) Expression patterns of Ca^2+^ genes during the temporal progression towards the respective lineage endpoints reveal large-scale, lineage-specific changes. (**F**) Venn diagrams illustrating the Ca^2+^ gene overlaps between the three lineages. (**G**) Principal tree graph calculated using scFates captures the bifurcation of Ca^2+^ signaling genes in the dentate gyrus (DG) and olfactory (OL) lineage branches (**H**) Heatmap of genes upregulated in the OL-branch (top) and DG-branch (middle) pre- and post-bifurcation (**I**) TF-Ca^2+^ gene networks showing the lineage-specific TFs that directly bind lineage-specific Ca^2+^ genes for each of the three lineages.

To further examine the regulatory links between Ca^2+^ signaling and the DG and OL lineages, we correlated TFs to the Ca^2+^-defined trajectories (Fig. S7F) and assessed whether Ca^2+^ genes were among the inferred targets of the top lineage-correlating TFs. Our analysis captured several TFs previously reported as regulators of neurogenesis within their specific lineages, such as *Rfx3, Bcl11b*, and *Prox1* in the DG (*21–23*), *Meis2* and *Pbx1* in the OL bulb (*24, 25*), and *Etv1* in the CB (*26*), indicating that the Ca^2+^ genes accurately reconstructed the lineage trajectories. Additionally, the TFs converged on Ca^2+^ genes that exhibited lineage-specificity, such as *Ncald*, *Kcnip1*, and *Camk4* (Fig. 3I). Our findings collectively suggest that the expression of Ca^2+^ signaling genes becomes increasingly lineage-specific during postnatal development, likely driven by the direct upregulation of distinct sets of Ca^2+^ genes by key lineage-determining TFs.

### Regional and functional heterogeneity of the Ca^2+^ signaling transcriptome during human embryonic neurogenesis

The lineage-specific expression of Ca^2+^ genes during postnatal neurogenesis inevitably raises the question of whether these genes exhibit similar trends during embryonic development. We therefore analyzed subsets of a scRNA-seq brain dataset generated from human embryos at 8 to 10 post-conception weeks (PCW) (*27*). We re-annotated the cells, guided by Braun, Danan-Gotthold et al. (*28*) (Fig. S8), and employed a similar Ca^2+^-module-based workflow as for the adult mouse nervous system, with the addition of removing cell cycle-driven modules (Fig. S9A-C; Methods). We found that the cells were predominantly organized along both a vertical glioblast-to-neuronal axis and a horizontal excitatory-to-inhibitory axis, with an apparent regional separation within both axes (Fig. 4A). The Ca^2+^ states were largely congruent with the cell states, capturing most of the heterogeneity in the dataset (Fig. S9D). The distribution of cell-module scores indicated that most neuronal modules were cell state-specific, with, for example, modules 3, 7, 9, and 14 marking the cerebellar inhibitory neurons, neocortical pyramidal neurons, cerebellar granule neurons, and diencephalic inhibitory neurons, respectively (Fig. 4B). Similarly, within the early progenitors, the Ca^2+^ modules were enriched in a region-dependent manner (Fig. 4C). The cell-gene scores for the VGCCs, CAMKs, CRAC-channels, and ATPases showed that individual members were differentially enriched across the cell states (Fig. 4D). Within the VGCCs, for example, *CACNA1H* was specific for the *EMX1+* lineage, mainly in the immature populations, while *CACNA1E* and *CACNA1A* were specific for the more mature pyramidal neurons. In contrast, *CACNA1C*, *CACNA1D*, and *CACNA1G* were predominantly high among the inhibitory neurons.

**Fig. 4.**
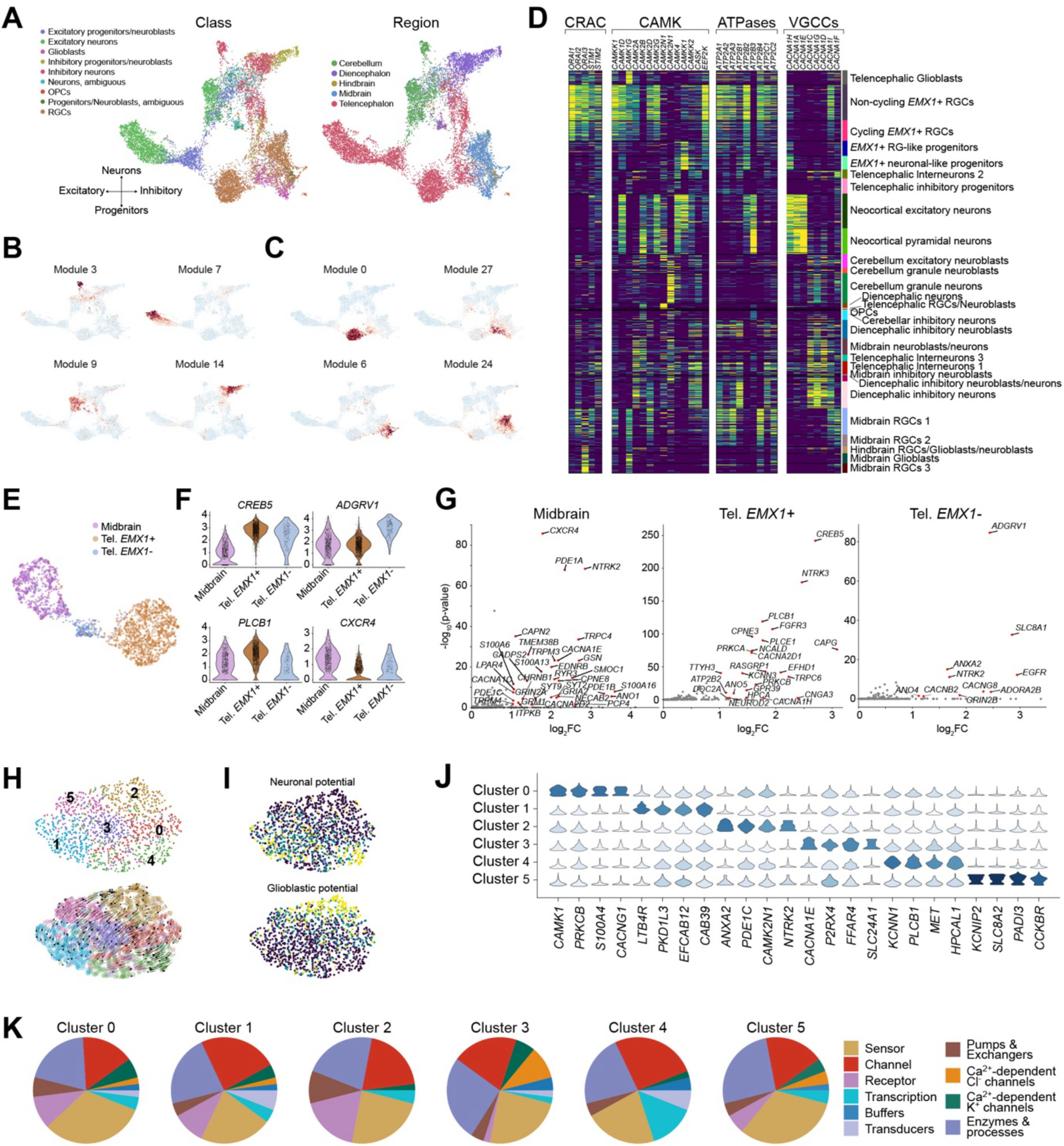
Transcriptional Ca^2+^ dynamics across early human brain development. (**A**) A Ca^2+^ based UMAP embedding of data from Van Bruggen et al. (*27*), colored by class (left) and region (right) (**B**) Feature plots illustrating examples of neuronal cell state-specific Ca^2+^ modules (**C**) Feature plots illustrating the regional specificity of Ca^2+^ modules within progenitors. (**D**) Heatmap showing the differential enrichment of individual members of canonical Ca^2+^ signaling. From left to right: Ca^2+^ release-activated channels (CRAC), Ca^2+^-/Calmodulin kinases (CAMK), ATPases; Voltage-gated Ca^2+^ channels (VGCCs). (**E**) Ca^2+^ based UMAP plot of RGCs found in the dataset, illustrating the clear regional separation into midbrain, dorsal telencephalon (*EMX1*+) and ventral telencephalon (*EMX1*-). (**F**) Violin plots showing examples of differentially expressed Ca^2+^ genes across the three RGC regions. (**G**) Scatter plots of differentially expressed Ca^2+^ genes across the three regions. (**H**) UMAP embedding of *EMX1+* RGCs, colored by functional clusters (*top*) with velocity vectors overlayed (*bottom*). (**I**) UMAP plots showing the neuronal and glioblast potential across the *EMX1*+ RGCs. (**J**) Violin plots illustrating the differential enrichment of Ca^2+^ genes across the clusters. (**J**) Pie charts of the gene classes of the top 50 genes per cluster.

Given the regional specificity of the Ca^2+^ modules within the early progenitor populations and the implication of Ca^2+^ signaling in driving RGC function (*8, 29*), we next sought to investigate regional differences in the expression of Ca^2+^ genes in RGCs. Although we annotated clusters as either “RGCs” or “Glioblasts”, there were recurring signs of intra-cluster variability within both populations (Fig. S10A). Accordingly, we calculated RGC scores for all cells based on the expressions of *HES1* and absence of *BCAN* and *NHLH1*, using these scores to identify RGCs (Fig. S10C, Methods). The RGCs were predominantly from the *EMX1+* dorsal telencephalon which eventually develops into the neocortex, the *EMX1-* ventral telencephalon and the midbrain (Fig. 4E). Interestingly, distinct sets of Ca^2+^ genes were upregulated in each developmental compartment (Fig. 4F-G) with, for example, *CREB5* marking the telencephalon, *ADGRV1* and *PLCB1* specifically marking the ventral and dorsal telencephalon, respectively, and *CXCR4* marking the midbrain.

Because RGCs in the neocortical ventricular zone are recognized for their spontaneous Ca^2+^ oscillations with varying frequencies and amplitudes during embryonic development (Fig. S10D) (*29*), we hypothesized that *EMX1*+ RGCs exhibit transcriptional variations in their Ca^2+^ signaling genes. By re-iterating our clustering strategy on the RGCs (Fig. S10E, Methods), we identified six distinct Ca^2+^- clusters within RGCs (Fig. 4H). Each of these clusters was enriched for different Ca^2+^ modules (Fig. S10F), and RNA velocity vectors indicated that the RGCs were organized along a maturity gradient from clusters 1 and 4 towards cluster 2 (Fig. 4H). Furthermore, computing the transition probabilities towards glioblasts or neurons based on the Ca^2+^ genes (Fig. S10G) indicated that clusters 2 and 4 exhibited the highest differentiation potential towards glioblasts and neurons, respectively (Fig. 4I). Notably, our analysis identified multiple cluster-specific Ca^2+^ genes, such as *ANXA2, LTB4R, KCNIP2, PRKCB, CACNA1E*, and *PLCB1* (Fig. 4J).

Next, we scored the Ca^2+^ genes for each cluster and plotted the distribution of the top 50, organized by their functional groups. Across all clusters, “Channels”, “Sensors”, and “Enzymes & Processes” were the dominant functional groups (Fig. 4K). Nevertheless, we found notable differences between the clusters. For example, comparing clusters 2 and 4, cluster 2 had a larger proportion of “Receptors” and “Pumps/Exchangers”, while cluster 4 had no “Receptors” but a high proportion of “Transcription” and “Transduction”.

Beyond the Ca^2+^ genes, differential expression analysis between clusters 2 and 4 revealed that genes expressed in late RGCs, such as *HOPX* and *MOXD1*, were upregulated in cluster 2, while genes expressed in early RGCs, including *NEUROG2* and *DLL1*, were upregulated in cluster 4 (Fig. S10H). This indicates that the two clusters consist of late and early RGCs, respectively, consistent with the divergence of differentiation trajectories predicted by the Ca^2+^ genes. Furthermore, it highlights the functional significance of the transcriptional variations in Ca^2+^ signaling genes among RGCs and offers another example of how differences in the expression of Ca^2+^ genes provide clues to functional distinctions.

### Delineation of Ca^2+^ dynamics during human neocortical development

Despite numerous studies highlighting the role of Ca^2+^ signaling in the regulation of neocortical development, the specific Ca^2+^ genes involved and how Ca^2+^ genes are remodeled along neurogenesis to adapt functionally remain unresolved. To address these questions, we analyzed the Ca^2+^ dynamics in the *EMX1*+ lineage. We found that the Ca^2+^-states were well-aligned to the cell states and that the Ca^2+^ genes accurately reconstructed the developmental trajectory from RGCs to pyramidal neurons, through the intermediate progenitor (IP) populations (Fig. 5A), with the RGCs as initial states and pyramidal neurons as terminal states (Fig. S11A). Notably, this could not be attributed to differences in the number of expressed Ca^2+^ genes or the total Ca^2+^ gene counts across cell states (Fig. S11B). Instead, our results suggested that the trajectory was driven by the up- and down-regulation of different Ca^2+^ genes over time (Fig. 5B-C). To test this hypothesis, we clustered the Ca^2+^ genes that were significantly associated with the neocortical trajectory based on their pseudotemporal kinetics (Methods). We found a total of 106 Ca^2+^ genes that grouped into 23 gene programs along the developmental trajectory (Fig. 5D-E; Fig. S11C). Most gene programs consisted of genes that were either expressed in RGCs and thereafter downregulated, or genes that were upregulated during development, peaking at different stages. Notably, some of the early gene programs remained temporally active in the earlier IPs, while some of the neuronal gene programs had already been switched on in the later IPs. Furthermore, certain gene programs were either transiently upregulated or downregulated, and a few genes, including *CREB5, KCNIP4*, and *CACNA2D1*, each formed their own group.

**Fig. 5.**
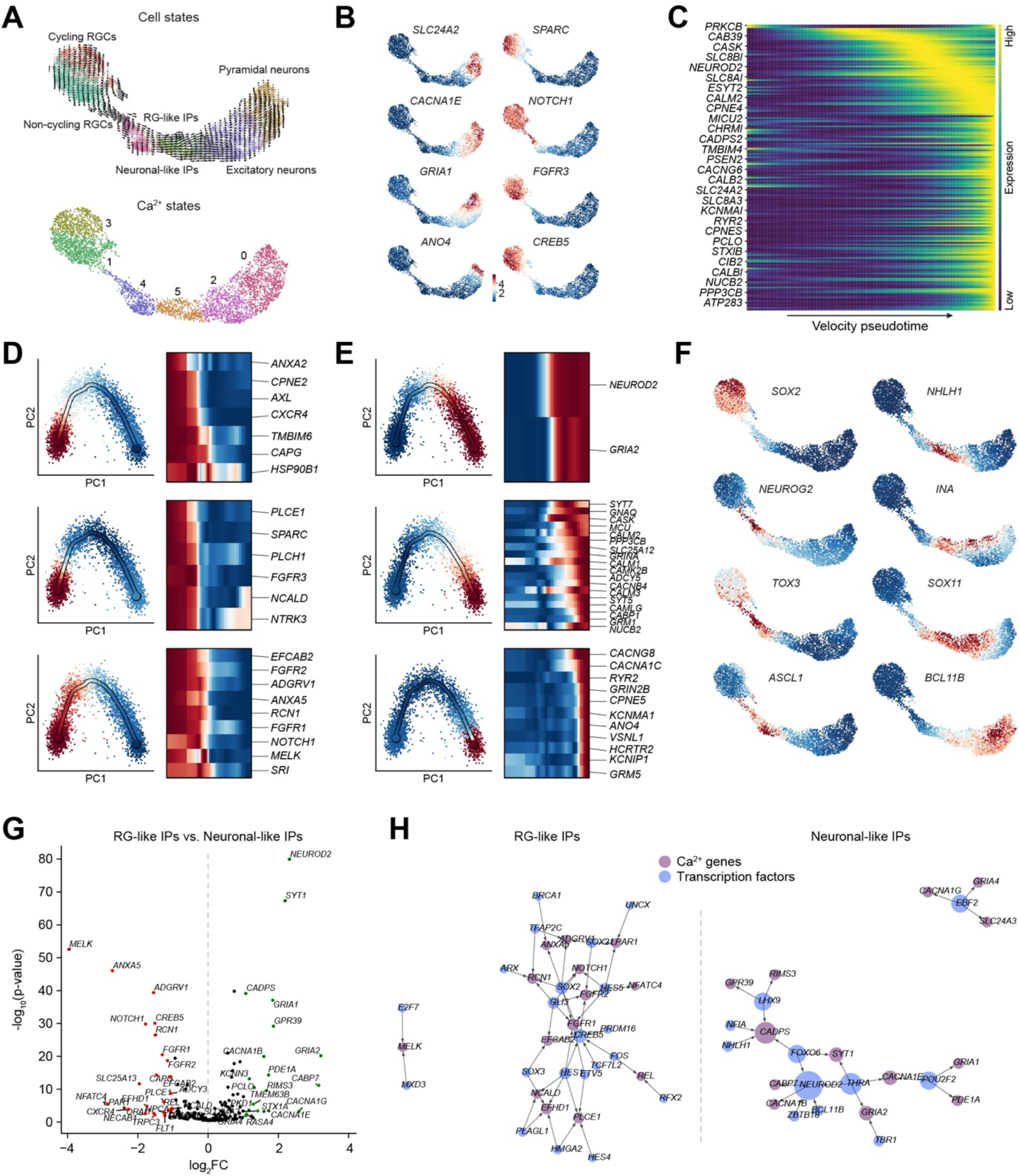
Delineation of Ca^2+^ dynamics during human neocortical development. (**A**) A Ca^2+^ based UMAP plot illustrating the trajectory of the entire *EMX1*+ lineage from Van Bruggen et al. (*27*), calculated using Ca^2+^ genes by cell state with velocity vectors overlayed (top) and Ca^2+^-state (bottom). (**B**) Feature plots illustrating examples of Ca^2+^ genes associated with the initial and terminal states. (**C**) Heatmap showing large-scale temporal changes in the expression of Ca^2+^ genes along the pseudotime trajectory. (**D-E**) Examples of the 23 defined Ca^2+^ gene programs along neocortical development using scFates, illustrating early (**D**) and late (**E**) programs. (**F**) Feature plots illustrating the expression of key regulatory TFs related to proliferation (left column) and differentiation (right column) (**G**) Volcano plot of Ca^2+^ genes upregulated in the differentiating neuronal-like intermediate progenitors (IPs) (green) respectively proliferative RG-like IPs (red). (**H**) TF-Ca^2+^ gene networks showing differentially expressed TFs and their predicted targets within the RG-like and the neuronal-like IPs.

Examining more closely the Ca^2+^ driven differences between the IPs, we found that the boundaries between the Ca^2+^-states aligned well with those from the whole genome and matched the switching of transcription factors that occurs as IPs transition from a proliferative, self-renewal state expressing *SOX2, NEUROG2*, *TOX3*, and *ASCL1,* to a differentiating state expressing *NHLH1, INA, SOX11,* and *BCL11B* (Fig. 5F), as reported elsewhere (*28, 30*). Moreover, differential expression analysis revealed that earlier IPs expressed RG-like Ca^2+^ genes while the latter IPs expressed neuronal-like Ca^2+^ genes (Fig. 5G). Many of the Ca^2+^ genes appeared to be regulated by *SOX2* and *NHLH1*, among others (Fig. 5H), implicating a role of Ca^2+^ signaling in regulating the phenotypic switch from proliferation towards differentiation.

### Conservation of Ca^2+^ dynamics during human neocortical development

To assess the evolutionary conservation of the Ca^2+^ signaling transcriptome during cortical development, we mapped cell types restricted to only Ca^2+^ genes (*31*) onto the mouse neocortical dataset of Telley et al. (*32*), spanning from embryonic day 12 (E12) to postnatal day 0 (P0) (Fig. 6A; Fig. S12A; Table S6). A significant portion (87 - 94%) of the mouse apical progenitors were classified as RGCs. Additionally, a majority (87 - 100%) of the 4-day-old neurons were categorized into one of two neuronal classes, indicating that the mapping was accurate. Notably, we observed a significant overlap between the 1-day-old neurons and IP cohorts. Among the 1-day-old neurons, for example, approximately half (46 - 56%) were predicted to be IPs, primarily neuronal-like. Although this contrasted with the original annotations, we found that these cells retained a higher expression of *Eomes* (Fig. S12B), consistent with the Ca^2+^-driven mapping.

**Fig. 6.**
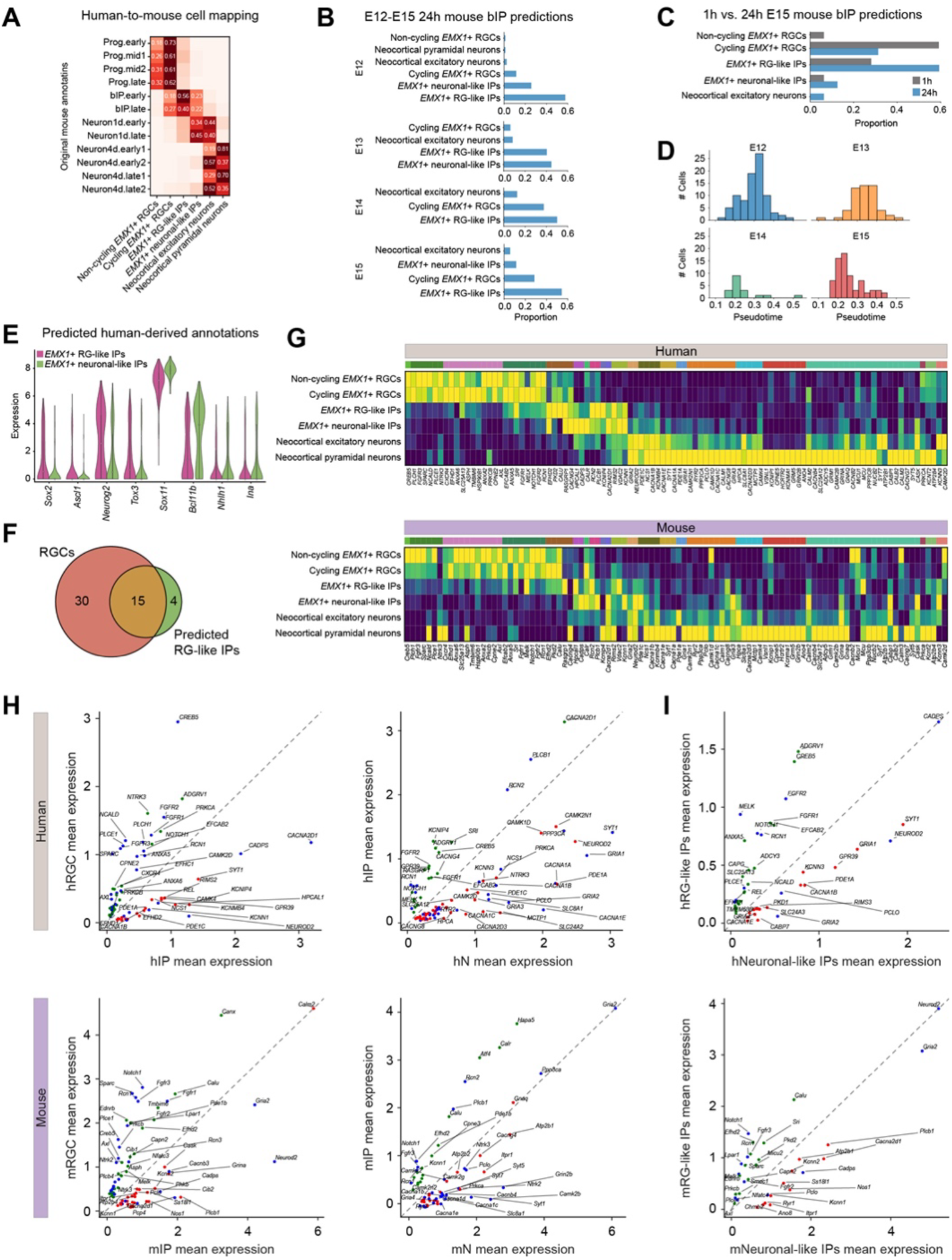
Human-defined Ca^2+^ dynamics are conserved in mouse neocortical development. (**A**) Confusion matrix illustrating the distribution of predictions across the cell types found in Telley et al. (*32*) using scArches, with each row summing to 1. Only values >0.1 are shown. (**B**) Bar plots illustrating the predicted distributions of 24h old mouse IPs, separated by birth date from E12-15. (**C**) Bar plots illustrating the predicted distributions of 1h vs. 24h E15 mouse IPs. (**D**) Histograms of pseudotimes of mouse IPs, separated by birth date, highlighting a left-shift with increasing birth date. (**E**) Violin plots of mouse IPs, grouped by their predicted human-defined cell states, show non-Ca^2+^ TF expressions consistent with the predictions. (**F**) Venn diagram of differentially expressed genes in the mouse RGCs and predicted RG-like mouse IPs. (**G**) Matrix plot showing the expression of genes belonging to the neocortical gene programs in the human data (top) and mouse data (bottom), grouped by predicted label. Every row constitutes the mean expression of the cell population. Every column constitutes a gene. Genes are colored by the gene programs. (**H**) Scatter plots of RGCs versus IPs (left) and IPs versus neurons (right) in humans (top) and mice (bottom). Every point represents the mean expression within each population. Blue dots denote conserved Ca^2+^ genes and green dots denote significantly upregulated genes in the developmentally more immature population, while red denotes the opposite. Only differentially expressed Ca^2+^ genes are shown. (**I**) Same as (**H**), but for RG-like versus neuronal-like IPs.

Although the mouse IPs have previously been described as two distinct age-defined populations, we found that both groups exhibited a similar distribution in terms of their class assignment. However, stratifications by birth dates revealed that IPs born during E12-E13 and E14-E15 predominantly resembled neuronal-like IPs and RG-like IPs, respectively (Fig. 6B; Fig. S12C). Additionally, comparisons of 1-hour-old and 24-hour-old IPs within the E15 population revealed that a greater proportion of the younger cells were classified as RGCs (65% compared to 29%), while most of the older cells tended to be categorized as RG-like IPs (28% compared to 54%) (Fig. 6C). Re-analysis of the mouse IPs indicated that they were organized along a pseudotime gradient that inversely correlated with their birthdate (Fig. 6D), and that the predicted neuronal-like IPs had a higher expression of neuronal transcription factors such as *Neurod2*, *Nhlh1*, and *Bcl11b*, whereas the predicted RG-like cells had a higher expression of *Sox2, Ascl1*, and *Hes1* (Fig. 6E; Fig. S12D). Supporting their RG-like state, the predicted mouse RG-like IPs had an upregulation of 19 Ca^2+^ genes, 15 of which were also upregulated in the mouse RGCs (Fig. 6F; Fig. 6H, including *Sparc, Notch1*, *Fgfr1, Fgfr2, Lpar1*, and *Rcn1* (Fig. S12E). Moreover, differentially expressed Ca^2+^ genes in the RG-like IPs overlapped with the mouse RGCs (Fig. 6F), further supporting the accuracy of the Ca^2+^-driven mapping.

To directly assess the conservation of the Ca^2+^ genes, we compared the expression of the identified neocortical developmental Ca^2+^ gene programs across human and mouse. We found a strong consensus between the species, suggesting that these gene programs were largely conserved, although a few genes showed contrasting expression patterns (Fig. 6G). Furthermore, in our analysis of differentially expressed Ca^2+^ genes across each sequential developmental step, we consistently observed that the most enriched genes, including *CREB5* and *NCALD* for RGCs, *CACNA2D1* and *CADPS* for IPs, and *CACNA1A* and *PPP3CA* for neurons, were conserved (Fig. 6H), although the degree of conservation varied among cell types (Fig. S12F). Collectively, these findings highlight that Ca^2+^ genes not only capture nuanced variations in cell states but also demonstrate a significant degree of evolutionary conservation of Ca^2+^ signaling in neocortical development, further emphasizing the role of specific Ca^2+^ signaling pathways in deterministically controlling cell states.

## Discussion

Here, we have comprehensively examined the Ca^2+^ signaling transcriptome in the entire adult mouse nervous system and early human brain development. Despite Ca^2+^ being a universal intracellular signaling mediator that is involved in regulating essentially all neuronal functions, our results reveal that genes driving Ca^2+^ signaling are heterogeneously and dynamically expressed across the brain with different neuronal types expressing distinct combinations of the Ca^2+^ signaling toolkit that reflect the functionality of the cell. This was, for example, evidenced by the neocortical layer II-IV neurons separating into Ca^2+^-states related to differences in experience-dependent remodeling, the RGCs forming functional clusters that were predictive of their glial/neuronal potential and the separation of IPs that was attributable to the phenotypic switch from proliferation to differentiation. Although our study is the first to specifically study the diversity of the Ca^2+^ signaling transcriptome across the brain, a survey of recent scRNA-sequencing studies focused on profiling neuronal cell types across different timepoints and regions revealed a recurrence of Ca^2+^ genes used as population-defining transcriptional markers, such as *Cacna2d1* in mouse interneurons (*33*), *CALB1* in the human midbrain (*34*), *PVALB* specifically in human midbrain glutamatergic neurons (*11*), and *CACNB2* and *CACNG3* for OPCs in early human development (*28*). These studies are consistent with our findings and support the hypothesis that the diversity of Ca^2+^ signaling has previously been underestimated.

Naturally, the heterogeneity of Ca^2+^ signaling in neurons raises the question of whether it is a function of development, whether there may exist population-specific environmental factors that influence the transcriptional Ca^2+^ state of neurons, or whether they reflect underlying neural circuitry. The ability of Ca^2+^ genes to reconstruct and differentiate between developmental trajectories using lineage-specific combinatorial patterns of Ca^2+^ gene programs illustrates that the divergence in Ca^2+^-states already begins to be specified during the progenitor stages, and thus that neurons are likely inherently primed to express specific subsets of Ca^2+^ signaling genes. This is further supported by the large-scale heterogeneity observable in early human brain development, where cells ranging from RGCs to neurons express region- and lineage-specific Ca^2+^ genes. With previous reports implicating Ca^2+^ signaling in regulating neurogenesis across many regions (*9, 35*), this transcriptional diversity suggests that the Ca^2+^-driven mechanisms are likely lineage-specific. Our results indicate that they may be directly activated by key regulatory TFs. Simultaneously, the clear regional patterning in Ca^2+^ signaling genes observable in progenitors as well as neurons suggest that environmental factors influence the transcriptional output and may potentially reflect the interactions between Ca^2+^ signaling, morphogens and transcription factors.

Within neocortical development, many Ca^2+^ genes are highly dynamic and strictly regulated, thereby containing a tremendous amount of information, as seen by the accurate re-construction of the lineage, and capturing key differences within RGCs as well as IPs. Our findings implicate numerous previously unexplored genes, including *CREB5*, *NCALD, KCNIP4*, and *CACNA2D1*, that seem to be evolutionary conserved; these are potential candidates for further investigation into their roles in regulation during neurogenesis. Together, our findings are in line with previous studies showing the critical role of Ca^2+^ signaling in regulating various aspects of neurogenesis, including cell proliferation, differentiation, and migration (*29, 35*). However, our study adds a new layer of understanding by elucidating the lineage-specific dynamics of Ca^2+^ signaling, which has significant implications for our insight into neocortical development.

Nevertheless, our study is not without its limitations, particularly in correlating our expression data with actual functional experiments. Current methodologies allow us to measure Ca^2+^ signaling only in a limited number of cells within thin brain sections. To overcome this, it is imperative to enhance our throughput capabilities, enabling the monitoring of Ca^2+^ signaling across a vast array of neurons in three dimensions. Additionally, a more thorough examination of distinct Ca^2+^ signaling pathways within specific cell types could have been performed to unveil new genes critical to certain cellular functions. However, the focus of this study was to conduct a comprehensive analysis of the entire Ca^2+^ signaling transcriptome across the whole brain, a task that, to our knowledge, has never before been accomplished in the field of Ca^2+^ signaling.

In conclusion, our study reveals a substantial diversity of Ca^2+^ signaling transcriptome across the human and mouse brain, and shows that the differential output of Ca^2+^ signals is not only driven by the spatiotemporal dynamics of the Ca^2+^ signal, but points to the existence of cell state-specific functional priming that shapes and restricts the potential outputs, as evidenced by the ability of Ca^2+^ genes to capture fine-grained differences despite only totaling hundreds of genes. Accordingly, Ca^2+^ signaling may more accurately be viewed as a dynamic signaling system that is tailored to specific neuronal identities and functions.

## Materials and Methods

### The Ca^2+^ signaling gene list

The Gene Ontology Consortium annotations were searched using AmiGO 2 to find genes associated with Ca^2+^ signaling using the keyword “Calcium” and restricted to the organisms “Homo sapiens” and “Mus musculus”. Annotations with the evidence code equivalent to only “automatic assertion” or “with no evidence using manual assertion” were excluded, yielding 1,515 genes. The GO compiled list was merged with the 1,670 genes found in the Calcium Gene Database (CaGeDB), our previously developed online database that maps genes related to Ca^2+^ signaling and their associated diseases (*14*), totaling ∼2,000 genes, followed by checking for the inclusion of canonical Ca^2+^ signaling genes. The list was finally manually curated by surveying the genes against UniProt, GeneCards, and published literature to exclude genes with unclear links to Ca^2+^ signaling, yielding a final list of 604 Ca^2+^ signaling genes. The genes were subsequently annotated and grouped into 46 functional groups, each containing at least 5 genes.

### Characterization of Ca^2+^-states across the entire adult mouse nervous system

The raw Unique Molecular Identifier (UMI) counts (spliced and unspliced) and the cell metadata were provided directly by the authors (*15*). Non-neuronal cells were removed, totaling a final set of 73,603 neurons classified into 214 cell types. The total UMI count matrix (i.e., sum of spliced and unspliced) across all Ca^2+^ signaling genes was decomposed using the *scHPF* package (*36*) with default settings, generating two matrices, representing cell respectively gene scores across the modules, *K*. *K* was set at 27, determined by running multiple iterations of the *scHPF* pipeline and manually tuning *K* such that K equaled the maximum number of modules where , where, on average, 4 modules captured a minimum of 70% of the cells’ scores (Fig. S1A-B). Mutual information scores were used to examine the redundancy of factors (Fig. S1C). Unless otherwise stated, all downstream analysis and visualization was done through *scanpy*. Cells were clustered on the cell-module scores by constructing a *k*-nearest neighbors (KNN) graph on the Jensen-Shannon distance (JSD) between every cell (*k* = 15) and neighbors were pruned by defining an information radius around every cell equal to the 85^th^ percentile of distances between neighboring cells in the KNN graph. The Leiden algorithm was used to generate clusters with the resolution set at 1.5. Clusters with <20 cells were deemed as outliers and removed, yielding 33 initial clusters. To ensure that the clusters used in the downstream analysis were robust and represented stable, interpretable transcriptomic states, the clusters were further refined by 1) removing cells assigned to a cluster if <10% of the cells or <15 cells belonging to that cell type were found there, and 2) merging clusters with <250 cells into a neighboring cluster based on a dendrogram constructed on high-dimensional UMAP coordinates (Fig. S1D). This yielded a final set of 63,507 cells, clustered into 28 different Ca^2+^-states. For the UMAP, the kNN-graph was re-computed using the JSD with *k* = 50 without radius pruning.

### Comparisons of scHPF performances between Ca^2+^ genes and random genes

A subset of 20% of the dataset was used for assessment due to computational efficiency. Next, the genes were pre-filtered such that only genes expressed in at least 10 cells were included, and all Ca^2+^ genes were excluded to avoid overlap in the gene sets. A sampling strategy was devised to randomly sample genes while simultaneously attempting to control for differences in expression levels and spread between Ca^2+^ genes and random genes. For each Ca^2+^ gene, the mean was calculated, and an expression interval equal to a deviation of 10% was allowed. From all of the expressed genes, genes with a mean expression outside of this interval were filtered out, followed by randomly sampling a gene from the remaining ones. The deviation was added to ensure that each iteration of scHPF was running on a different set of random genes. To assess the performance of the iterations, the cells were labeled as their respective Ca^2+^-states and projected on UMAPs. The local inverse Simpson’s Index score (LISI) (*37*) was computed for each UMAP embedding through the package *harmonypy* and was used to compare the performance of each iteration. Briefly, for each cell, LISI measures the diversity in the local neighborhoods of cells where a score of 1 implies a homogenous neighborhood and higher scores imply a mixture of multiple neighborhoods.

### Building of random forest classifiers

The Python packages *scikit-learn* and *optuna* were used to build our random forest classifiers. Both the Ca^2+^ and random gene counts were first normalized and log-transformed. The classifiers were trained on 80% of the data, and the performances were evaluated by classifying the remaining held-out 20%. For each classifier, *Optuna* was used to optimize the classifiers, with ‘n_trials’ set to 100, and StratifiedShuffleSplit was used for cross-validation by generating multiple randomized folds (n_splits = 5). Separate classifiers were built for each of the two levels (Ca^2+^-states and cell types), totaling four. Similar to the above the random genes were matched to the Ca^2+^ genes for the classifiers by selecting genes with the lowest differences between the means compared to Ca^2+^ genes.

### Computation of cell-gene scores from cell-module scores and gene-module scores

To transform the cell-module and gene-module matrices back to a cell-gene matrix, *z*-scores were calculated from the cell-module scores and capped at a maximum value of 2.5. Next, the dot product of the z-scaled cell-module scores and the gene-module scores were calculated to yield a cell-gene matrix. The final cell scores were then obtained by z-normalizing the cell-gene matrix. For each Ca^2+^- state, the top Ca^2+^ genes were found by ranking the mean scores of all cells within the Ca^2+^-state. Unless otherwise stated, the top 50 genes were used.

### Enrichment of Ca^2+^ functional groups

The relative importance of the Ca^2+^ functional groups on the Ca^2+^-states was assessed by using *MetaNeighbor*, identical to the approach as described by Morarach (*38*), which. Simplistically, *MetaNeighbor* quantifies the ability of each functional group to discriminate between Ca^2+^ states. Functional groups with at least five genes were selected. The performance was expressed as the mean area under the receiver operator characteristic curve (AUROC) score. The normalized AUROC scores were calculated by dividing the AUROC scores by the mean AUROC scores for each gene set across the entire dataset.

### Relationship between Ca^2+^-state-1 and Ca^2+^-state-20 and TEGLU7, TEGLU8, and TEGLU12

To determine the relationship between *Ca^2+^-state-1* and *Ca^2+^-state-20* as well as their corresponding cell types (*TEGLU7, TEGLU8, TEGLU12*), conventional pre-processing was done through *scanpy*, including filtering lowly expressed genes, normalizing the counts and computing the most variable genes prior to running a principal component analysis (PCA). The linear discriminant analysis (LDA) was then calculated using the PCA cell loadings, with an incremental increase in the number of PCs.

### Calculations of gene regulatory networks

For the computation of enriched gene regulatory networks, the single-cell regulatory network inference and clustering (SCENIC) pipeline was run through the Python package *pySCENIC* (*39*). Genes expressed in <3 cells were first removed. The parameter “auc_threshold” was set at 0.01, and “nes_threshold” was set at 2.0, with the rest kept at default. For the developmental human and mouse data, the Ca^2+^ gene list was merged with the TF list provided in the package to capture potential Ca^2+^ genes with promoter binding activity. For the adult mouse data, the built-in regulon specificity score (RSS) function was used to find enriched regulons. For each of the two Ca^2+^-states, the top 20 TFs were selected. For each TF, the top 30 targets were selected, and only the differentially expressed were retained in the final comparison. The Spearman correlation was calculated across the entire dataset and included all of the predicted TFs. The TF-target gene networks were constructed using the networkx package.

### Gene set enrichment analysis

The *gget* package (https://github.com/pachterlab/gget) was used to perform the gene set enrichment analyses with databases Reactome_2022 and KEGG_2021_Mouse.

### Enrichment of synapse remodeling genes

The comparison of synapse remodeling-related genes across the cell types was computed using a list of genes adapted from Südhof (*17*). Differential expression analysis was done with the log foldchange threshold set at 0.5.

### Lineage inference analysis across the developmental subset of the adult mouse brain

For the lineage inference analysis across the developmental subset of the mouse brain data, the data was first subset to only include the developmental Ca^2+^-states. Next, *scHPF* was re-run with *K* instead tuned such that 3 modules on average captured 70% of the cells’ scores, yielding 18 modules. The KNN graph using the JSD was set at *k* = 100 and the Leiden resolution at 0.6. Clusters belonging to neither lineage were then removed. The lineage trajectory inferences were computed using the *CellRank* (*40*) and *scVelo* (*41*) packages. Standardized gene filtering and count matrix normalization were done (min_cells = 3) prior to the selection of Ca^2+^ genes, yielding 15,199 cells expressing 469 Ca^2+^ genes. Moments were calculated using ‘scvelo.pp.moments’ with n_pcs = 15 and n_neighbors = 30, and the velocity vectors were calculated with the mode set to ‘stochastic’. CellRank was then run with the default setting using a combined kernel weighting the “VelocityKernel” and “ConnectivityKernel” 80% to 20%. The lineage-determining TFs and Ca^2+^ genes were subsequently computed through the built-in function compute_lineage_drivers(method = ’perm_test’).

### Characterization of Ca^2+^-states across the early human development

To investigate the dynamics of Ca^2+^ signaling in early human brain development, data from Van Bruggen et al. (*27*) was used. Originally, the dataset was obtained using 10X Genomics with Chromium versions 2 and 3. To avoid batch corrections that may skew the results, only the version 3 cells were used, yielding a total of 12,990 cells from PCW 8.5 – 9. The cell annotations were done through comparisons to Braun, Danan-Gotthold et al. (*28*), and published literature. *scHPF* was run with the same criteria as for the adult mouse data, yielding 31 modules. To remove cell cycle-related modules, Pearson correlations were calculated between cell-module scores and the cell cycle markers *TOP2A, MKI67,* and *BIRC5*. Modules *11* and *16* were thus excluded from further analysis due to their high correlation coefficients, yielding 29 final Ca^2+^ modules. A clustering strategy similar to the adult mouse was subsequently used (neighbors = 15, information radius defined by the 85^th^ percentile), albeit with the Leiden resolution set at 2.5 and clusters with <20 cells removed.

### Analysis of human radial glial cells

To define radial glial cells (RGCs), the *scanpy* package function “scanpy.tl.score_genes” was separately run on the genes *HES1, BCAN,* and *NHLH1* across all cells. The scores were then binarized with the cutoff = 1. RGCs were then defined as cells with a *HES1* score of 1 and the rest at 0, totaling 2282 RGCs. Differential expression analysis was done by grouping the RGCs by developmental compartment (dorsal/ventral telencephalon and midbrain). The log foldchange threshold was set at 1. For the analysis of the heterogeneity within *EMX1+* RGCS, *scHPF* was run with the modules set at 50, and 3 cycling modules were removed. Clustering was done using the JSD distance (neighbors = 25, Leiden resolution = 0.7), yielding 6 clusters. Differential expression analysis between clusters 2 and 4 was done with the log foldchange threshold set at 0.5 and used as the basis for deducing early/late RGCs. To ensure that they were accurate, the velocity vectors were calculated using all genes.

### Lineage inference analysis across the human *EMX1+* lineage

First, the *EMX1+* RGCs, neuronal intermediates, and neurons were selected. Next, the cells were clustered based on Ca^2+^ genes, with the following parameters: number of neighbors = 15, distance = correlation, and Leiden resolution = 0.6. Following this, the lineage inference was done using the same settings for the mouse data outlined above. To relate RGCs to a potential glioblast or neuronal fate, the inference analysis pipeline was re-run with the inclusion of *EMX1+* glioblasts and manually setting the number of macrostates to 2 by using “g.compute_macrostates(n_states = 2)” prior to computing terminal states, ensuring that the absorption probabilities towards glioblasts would be calculated. The absorption probabilities towards neurons and glioblasts for RGCs were then *z-*score normalized and correlated to the clusters.

### Pseudotemporal and bifurcation analysis

The pseudotemporal analysis along the neocortical *EMX1+* lineage and the bifurcation analysis of the divergence towards the DG- and OL-lineages, respectively, were done using the *scFates* package (*42*). To ensure that the significant Ca^2+^ features would not be biased, the entire pipeline was first run using all genes. For the pseudotemporal trajectory analysis on the human data, the pipeline was run using the default settings. All non-Ca^2+^ genes were removed after running the “scfates.tl.test_association” function. The parameter “A_cut” was set to the 90^th^ percentile, resulting in a total of 106 significant Ca^2+^ genes that were subsequently fitted. To find gene programs, genes were clustered based on their fitted kinetics. First, the expression was normalized such that they summed to 1. Next, the Ca^2+^ genes were hierarchically clustered using the complete-linkage on the cosine distances, generating 23 gene clusters that were subsequently plotted using the *scFates* plotting interface. For the bifurcation analysis, a principal tree was generated using the “ppt” method on the UMAP-embedding. Next, the branch point was extracted. “A_cut” was set to 0.4 and the genes were calculated using “scfates.tl.test_fork” and “scfates.tl.branch_specific” with effect = 0.1 resulting in a total of 28 Ca^2+^ genes (18 DG; 10 OL).

### Ca^2+^ driven integration of human and mouse neocortex

The Python package *scArches* with the built-in model *scPoli* (*43*) was used to integrate the two datasets, using the human data as the reference and the mouse data as the query. It stands to reason that if the Ca^2+^ signaling transcriptome was generally conserved between human and mouse, mapping the mouse cells using the expression from human cells should be possible, assuming the existence of comparable cell populations across both datasets. Accordingly, all non-Ca^2+^ genes were removed, and only the shared Ca^2+^ genes were retained prior to the mapping. The integration was run using recommended settings based on the documentation tutorial, only changing n_epochs (=30) and pretraining_epochs (=24) within the function “scPoli.train”. The performance was evaluated by comparing the predicted human annotations to the original mouse annotations.

### Ca^2+^ imaging

Neocortical slices were prepared from mouse pups at embryonic day E12.5-18.5. Briefly, after brain extraction, 300-μm coronal slices were cut perpendicular to the pial surface with a vibratome (VT1000S; Leica) in ice-cold modified artificial cerebrospinal fluid (ACSF) containing the following: 125 mM NaCl, 25 mM NaHCO_3_, 1.25 mM NaH_2_PO_4_, 20 mM glucose, 2.5 mM KCl, 1 mM CaCl_2_, 8 mM MgCl_2_. After being allowed to rest for at least 1 h in oxygenated ACSF containing 125 mM NaCl, 25 mM NaHCO_3_, 1.25 mM NaH_2_PO_4_, 20 mM glucose, 5 mM KCl, 2 mM CaCl_2_, and 1 mM MgCl_2_ at room temperature (RT), slices were transferred to a custom-made oxygenated loading chamber, where they were bulk loaded with Fluo-4/AM (30 μM; Invitrogen) and 0.07% Pluronic F-127 (Invitrogen) at 37°C for 30 to 40 min. Slices were then allowed to rest and rinse in ACSF at RT for least 30 min before being placed into the acquisition chamber and imaged with a 2-photon laser-scanning microscope (2PLSM, Zeiss LSM 510 NLO; Carl Zeiss) equipped with a Coherent Chameleon Ultra II Titanium-sapphire pulsed laser (Coherent, Inc.) and a plan-apochromat 20× 1.0 NA water immersion objective lens (Carl Zeiss). Slices were held in place with a nylon mesh grid (Warner Instruments), and oxygenated ACSF (at RT) was perfused into the chamber continuously during the experiments. All images were acquired at a tissue depth of at least 30 μm. Fluo-4/AM was excited at 910 nm, and emission light was spectrally filtered by a 510- to 600-nm bandpass filter. A sequence of 16-bit single z-plane images was acquired at a 512 × 512-pixel resolution with a sampling frequency of 0.5 Hz (every 2 s) under the control of the Zeiss Zen software (Carl Zeiss). Custom in-house MATLAB scripts and ImageJ were used to analyze the data.

## Acknowledgments

We would like to thank Dr. David van Bruggen for valuable discussions throughout this study. A special thanks to Dr. Sten Linnarsson, Dr. Gonçalo Castelo-Branco, and Dr. Peter Lönneberg for providing access to data and high-performance cluster and Uhlén lab members for critical feedback.

## Funding

This work was supported by:

Swedish Research Council 2017-00815 (PU)

Swedish Research Council 2021-03108 (PU)

Swedish Research Council 2022-02185 (ES)

Swedish Brain Foundation FO2018-0209 (PU)

Swedish Brain Foundation FO2020-0199 (PU)

Swedish Cancer Society 19 0544 Pj (PU)

Swedish Cancer Society 19 0545 Us (PU)

Swedish Cancer Society 22 2454 Pj (PU)

Swedish Childhood Cancer Foundation PR2020-0124 (PU)

Swedish Childhood Cancer Foundation PR2022-0111 (PU)

Karolinska Institutet’s CSTP doctoral program (IAR)

## Author Contributions

Designed the study I.A.R., L.L., E.S., and P.U.

Performed and analyzed the Ca^2+^ experiments L.L. and I.D.E.

Analyzed the transcriptional data I.A.R.

Wrote the paper I.A.R. and P.U. Oversaw and conceived the study P.U.

## Competing interests

Authors declare that they have no competing interests.

## Data and materials availability

All data are available in the main text or the supplementary materials.

## Supplementary Materials

Materials and Methods

Figures S1-S12

Table S1-S6

References

## Notes

### Competing Interest Statement

The authors have declared no competing interest.

