## Supplementary Figures for "Single-Cell Transcriptomics Reveals the Molecular Logic Underlying Ca^2+^ Signaling Diversity in Human and Mouse Brain"

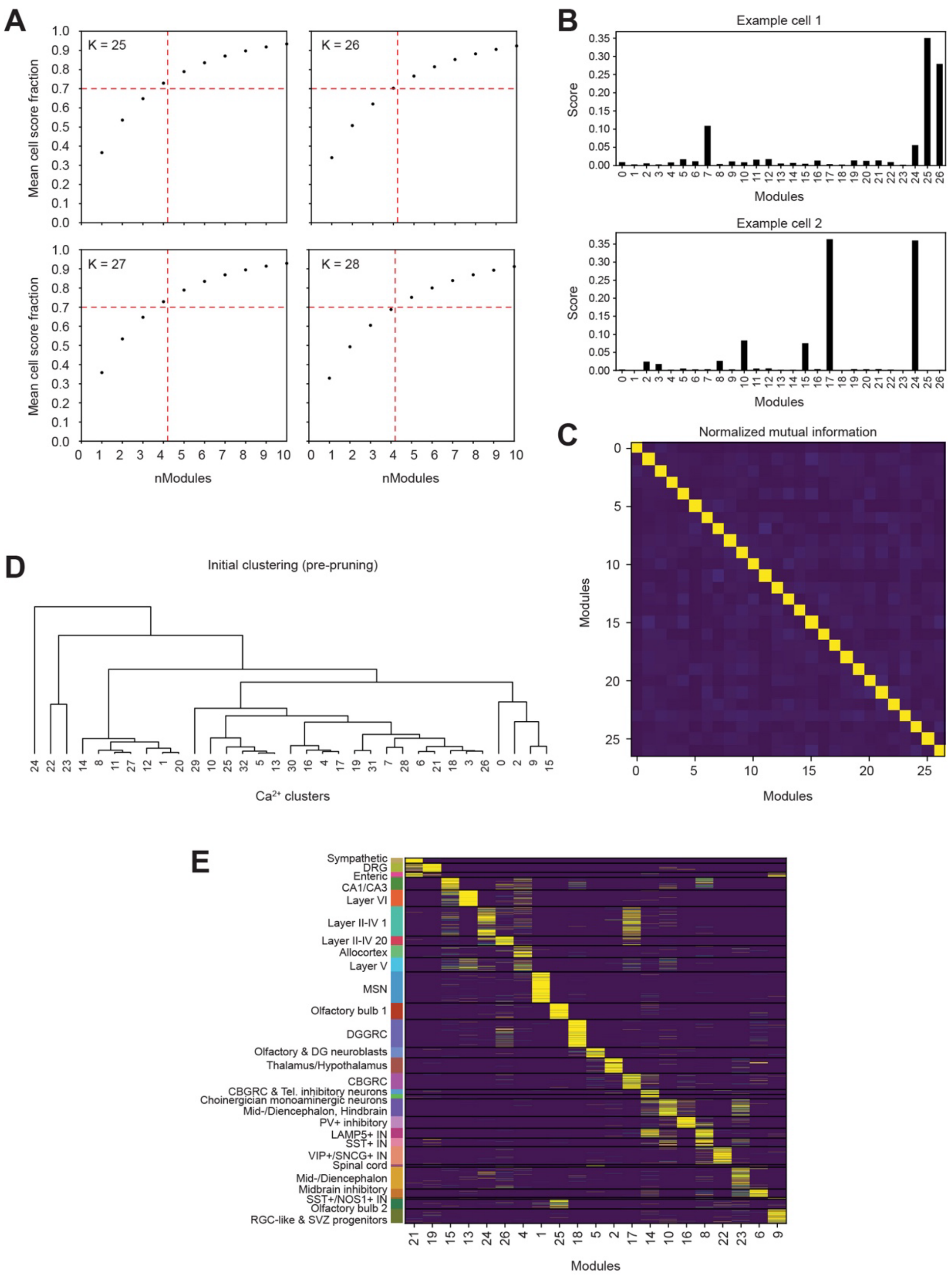

**Fig. S1. Overview of the HPF-based clustering strategy.** (**A**) Mean cell score fraction plotted against the number of modules. For the dataset from Zeisel et al. (*15*), scHPF was iteratively run with *K* manually tuned to the maximum value where four modules captured a mean of 70% of the cell scores. (**B**) Examples of cell-module scores for two randomly sampled cells, illustrating the differential enrichment of modules across the atlas. (**C**) Heatmap of the normalized mutual information scores between the modules. (**D**) Dendrogram of the initial 33 $Ca^{2+}$ clusters, pre-pruning and merging the small clusters according to the hierarchical similarities. (**E**) Heatmap illustrating the differential enrichment of modules across the 28 final $Ca^{2+}$-states.

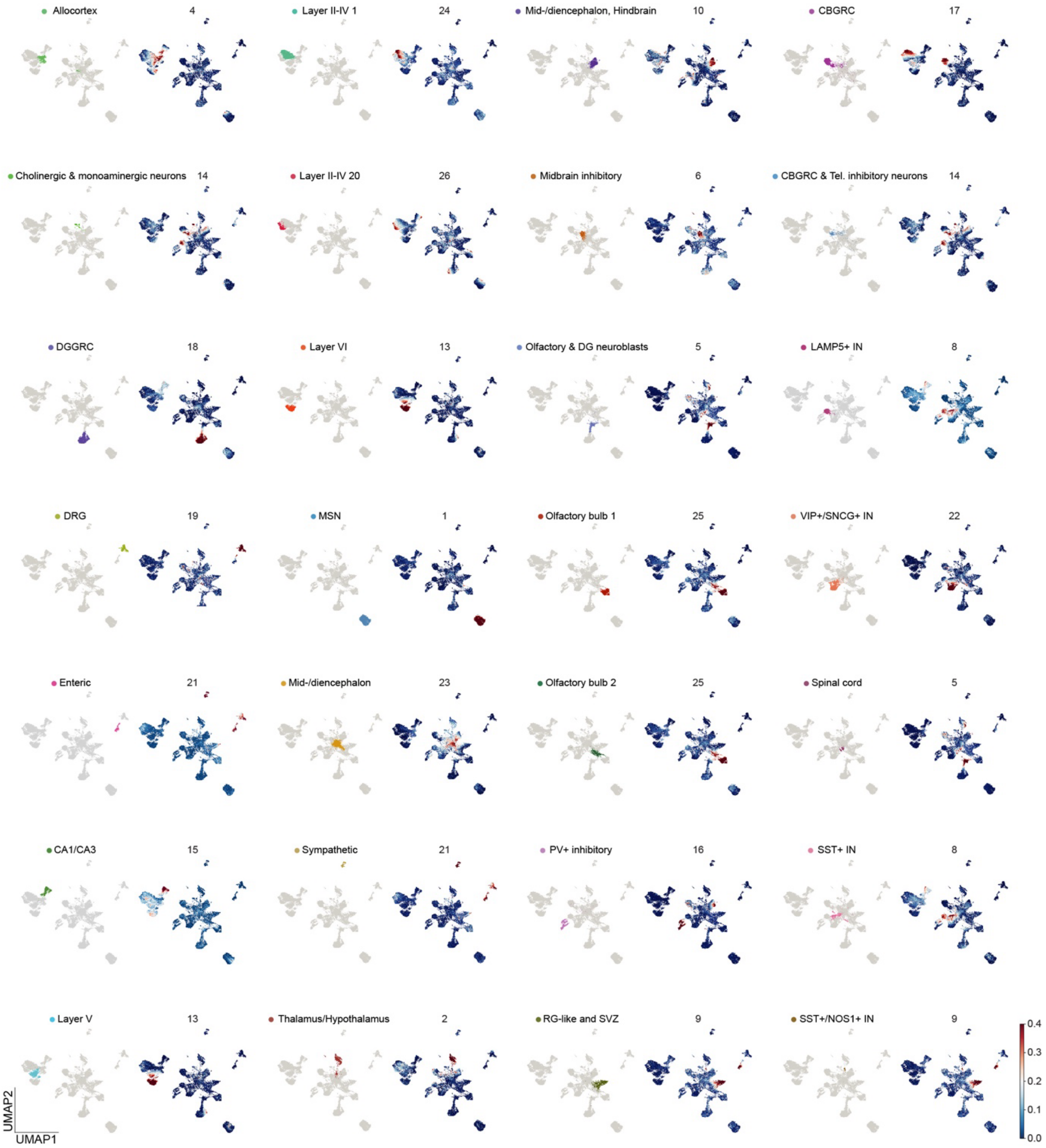

**Fig. S2. Illustrations of the 28 $Ca^{2+}$-states and their associated $Ca^{2+}$ modules.** UMAP plots of each individual $Ca^{2+}$-state (left) and their highest associated $Ca^{2+}$ module (right).

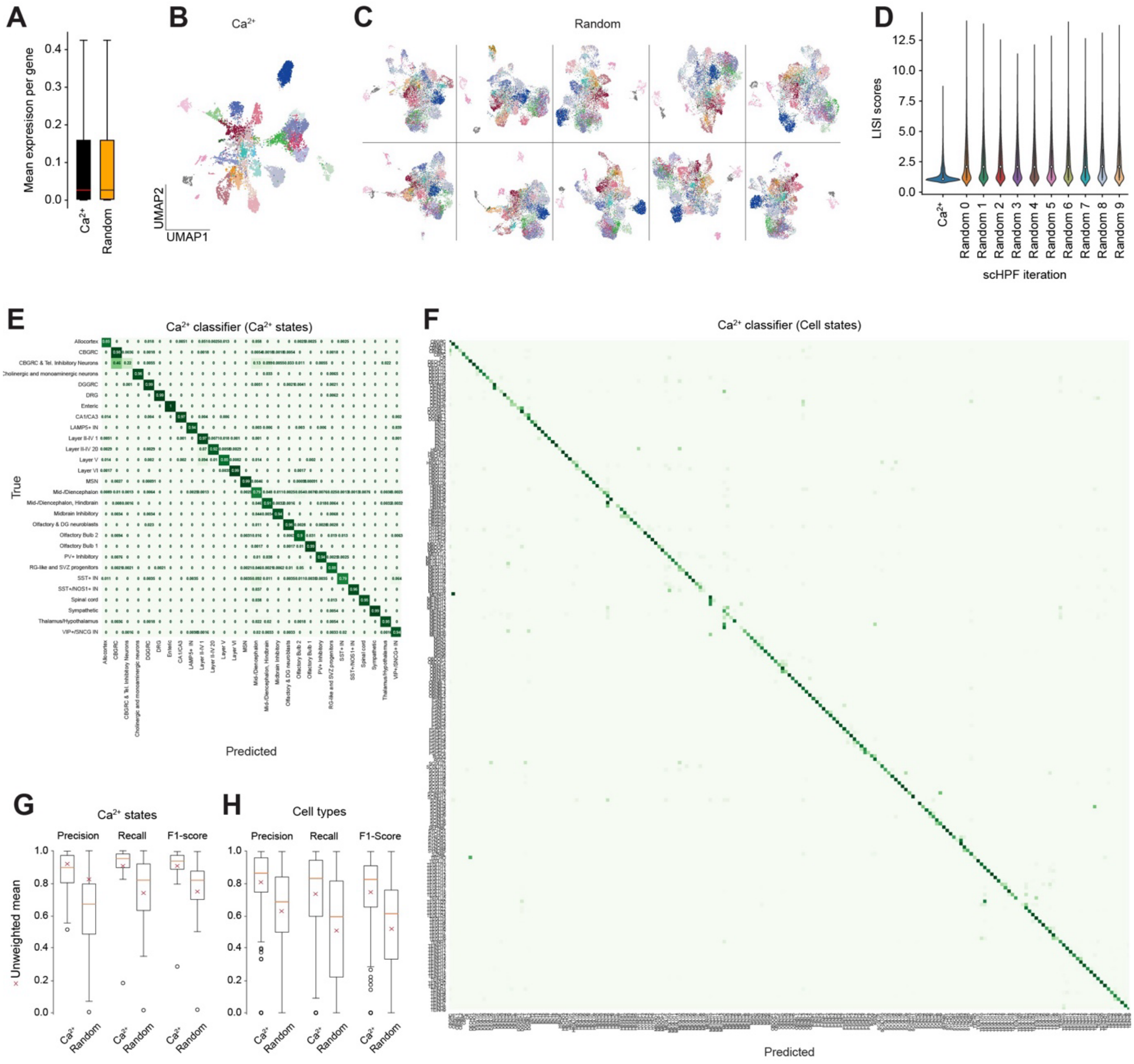

**Fig. S3. Comparisons between $Ca^{2+}$ and random genes.** (**A**) Boxplot showing the distribution of expressions of one iteration of matched random genes side-by-side with $Ca^{2+}$ genes, UMAP plot illustrating the divergent trajectories based on transcriptomic changes in $Ca^{2+}$ signaling genes. Arrows indicate $Ca^{2+}$-based velocity vectors. (**B**) UMAP plot based on scHPF of $Ca^{2+}$ genes calculated from a subset of the dataset from Zeisel et al. (*15*), colored by $Ca^{2+}$-states. (**C**) UMAP plots based on scHPF of ten different iterations of random genes, colored by $Ca^{2+}$-states. (**D**) LISI scores, representing neighbor heterogeneity (lower value indicates a more homogenous neighborhood), calculated on the $Ca^{2+}$ based UMAP and each of the ten iterations of random genes. (**E**) Confusion matrix showing the true and predicted $Ca^{2+}$-states using the $Ca^{2+}$ classifier. (**F**) Confusion matrix showing the true and predicted cell types using the $Ca^{2+}$ classifier. (**G**) Boxplots of the precision, recall and F1-scores of the $Ca^{2+}$ and random gene classifier on the level of $Ca^{2+}$-states. (**H**) Boxplots of the precision, recall and F1-scores of the $Ca^{2+}$ and random gene classifier on the level of cell types.

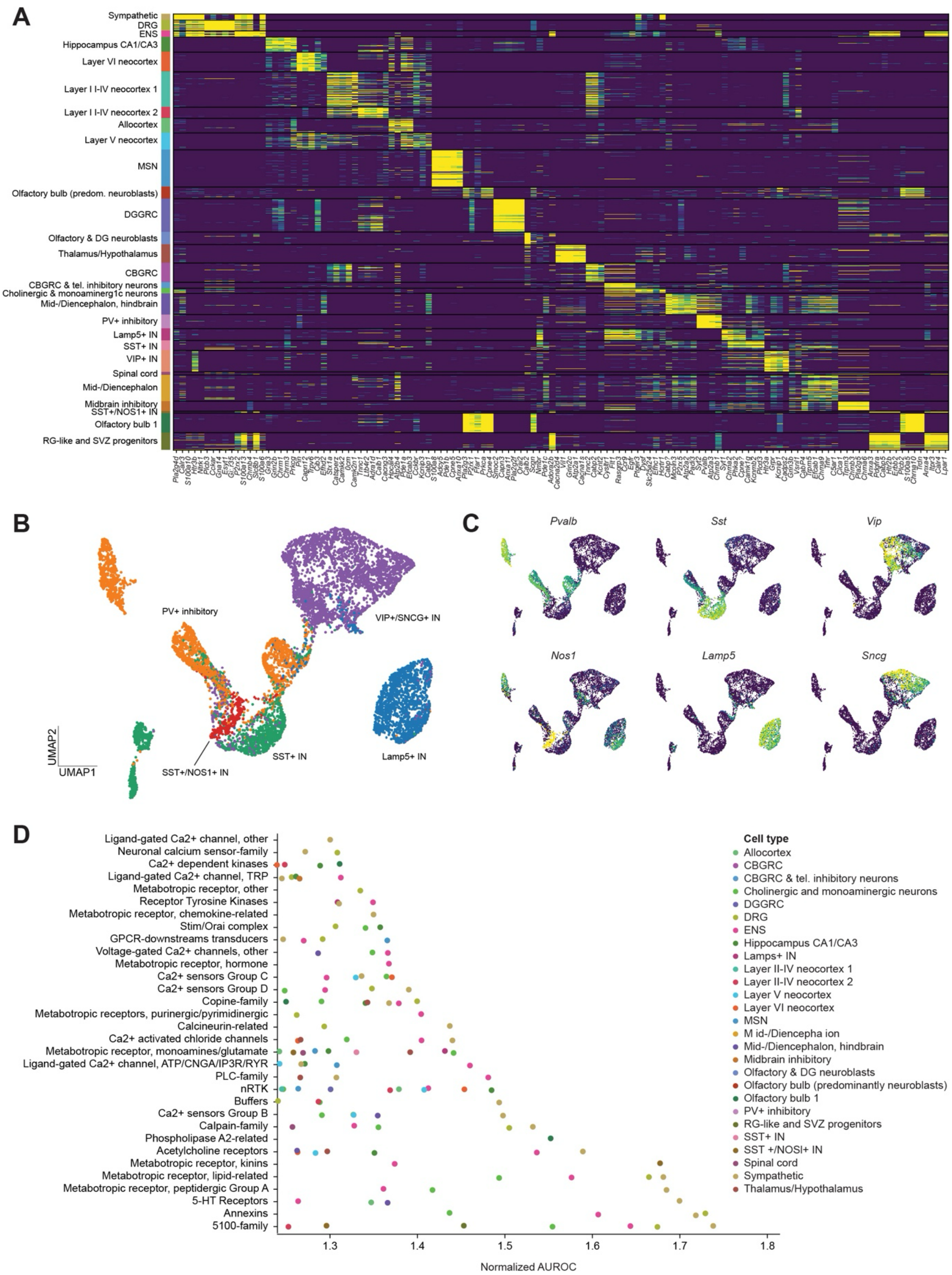

**Fig. S4. Further analysis of gene enrichments across** Zeisel et al. (*15*)**.** (**A**) Heatmap of cell-gene scores illustrating the top genes per $Ca^{2+}$-states across the entire dataset. (**B, C**) UMAP plots of interneurons colored by $Ca^{2+}$-states (**B**) and feature plots of marker genes used to identify sub-populations of cortical interneurons (**C**). (**D**) Normalized AUROC scores of heterogeneous $Ca^{2+}$ gene sets, visualizing $Ca^{2+}$-states with scores > 1.2.

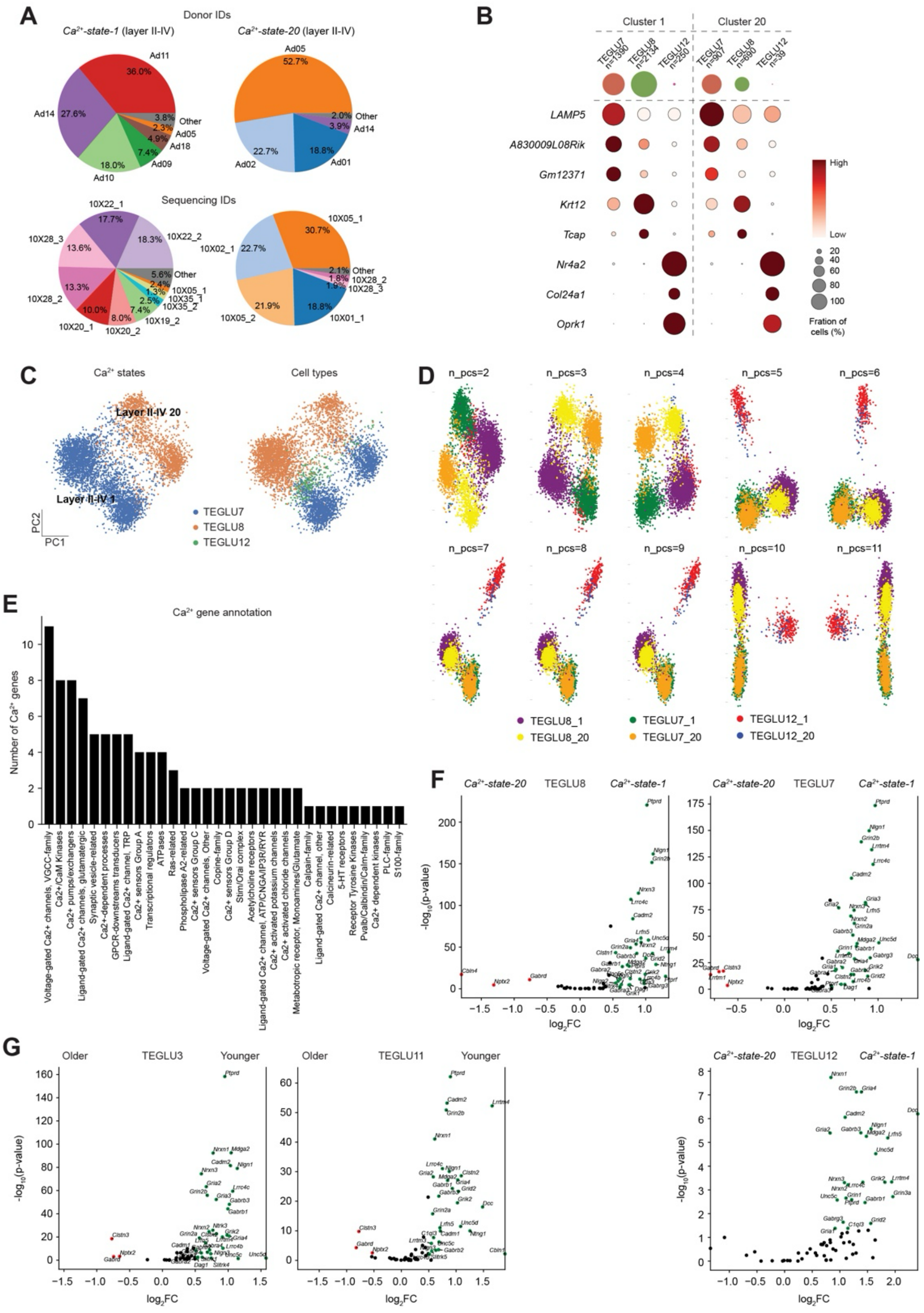

**Fig. S5. Further analysis of *Ca$^{2+}$-state-1* and *Ca$^{2+}$-state-20* in layer II-IV. Delineation of Ca$^{2+}$ dynamics during human neocortical development.** (**A**) Pie charts showing the distribution of Donor IDs (top) and experimental sequencing ID (bottom) of both Ca$^{2+}$-states. (**B**) Dot plot of the expressions of marker genes for each of the three cell types found in the layer II-IV Ca$^{2+}$-states, illustrating that they are shared across the Ca$^{2+}$-states. (**C**) Scatter plot of the first two principal components (PCs), colored by Ca$^{2+}$-state (left) and cell type (right). (**D**) Scatter plots of 2D Linear Discriminant Analysis, calculated on incremental increases of PC components, colored by cell type and Ca$^{2+}$-state. With the number of PCs <4, the separation is best explained by the Ca$^{2+}$-states. (**E**) Bar plot of differentially expressed Ca$^{2+}$ gens in *Ca$^{2+}$-state-1* versus *Ca$^{2+}$-state-20*, grouped by the gene set classification. (**F**) Volcano plots of synapse formation/remodeling-related genes, separated by cell types, across the two Ca$^{2+}$-states. (**G**) Volcano plots of the same genes across *TEGLU3* and *TEGLU11*, showing that similar age-related differences were observable in further cell types.

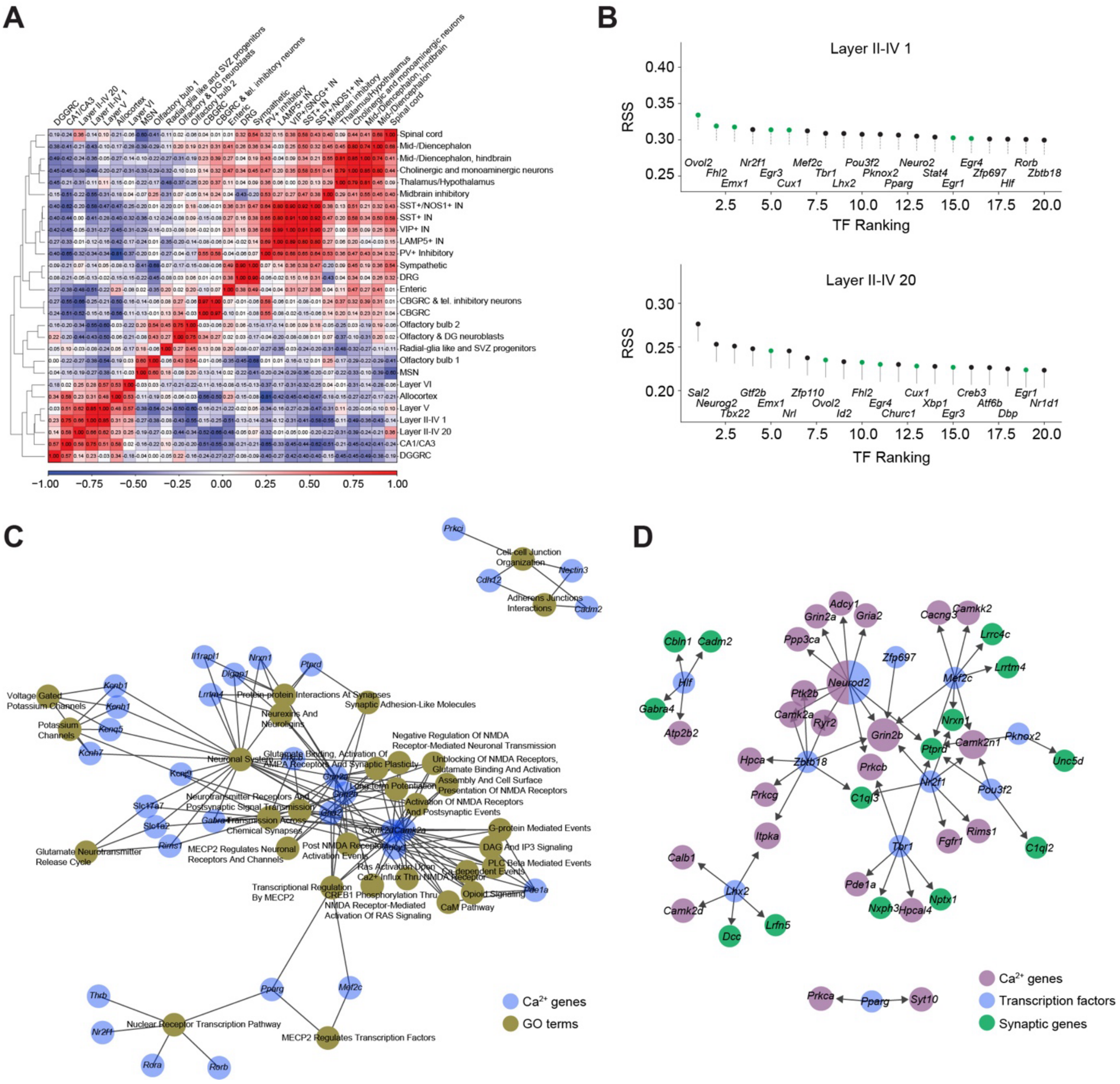

**Fig. S6. Gene regulatory network inference using pySCENIC.** (**A**) Correlation matrix calculated from predicted transcription factor (TF) activities across the entire Zeisel et al. (*15*) dataset (**B**) Top enriched TFs for the two layer II-IV neocortical states, calculated as the regulon specificity scores (RSS). Green highlights shared TFs. (**C**) Networks of enriched gene ontology terms and synapse-related genes constructed from the non-shared enriched TFs in *Ca²⁺-state-1.* (**D**) Networks of enriched TFs in *Ca²⁺-state-1* and their predicted synapse-related gene targets and $Ca^{2+}$ genes.

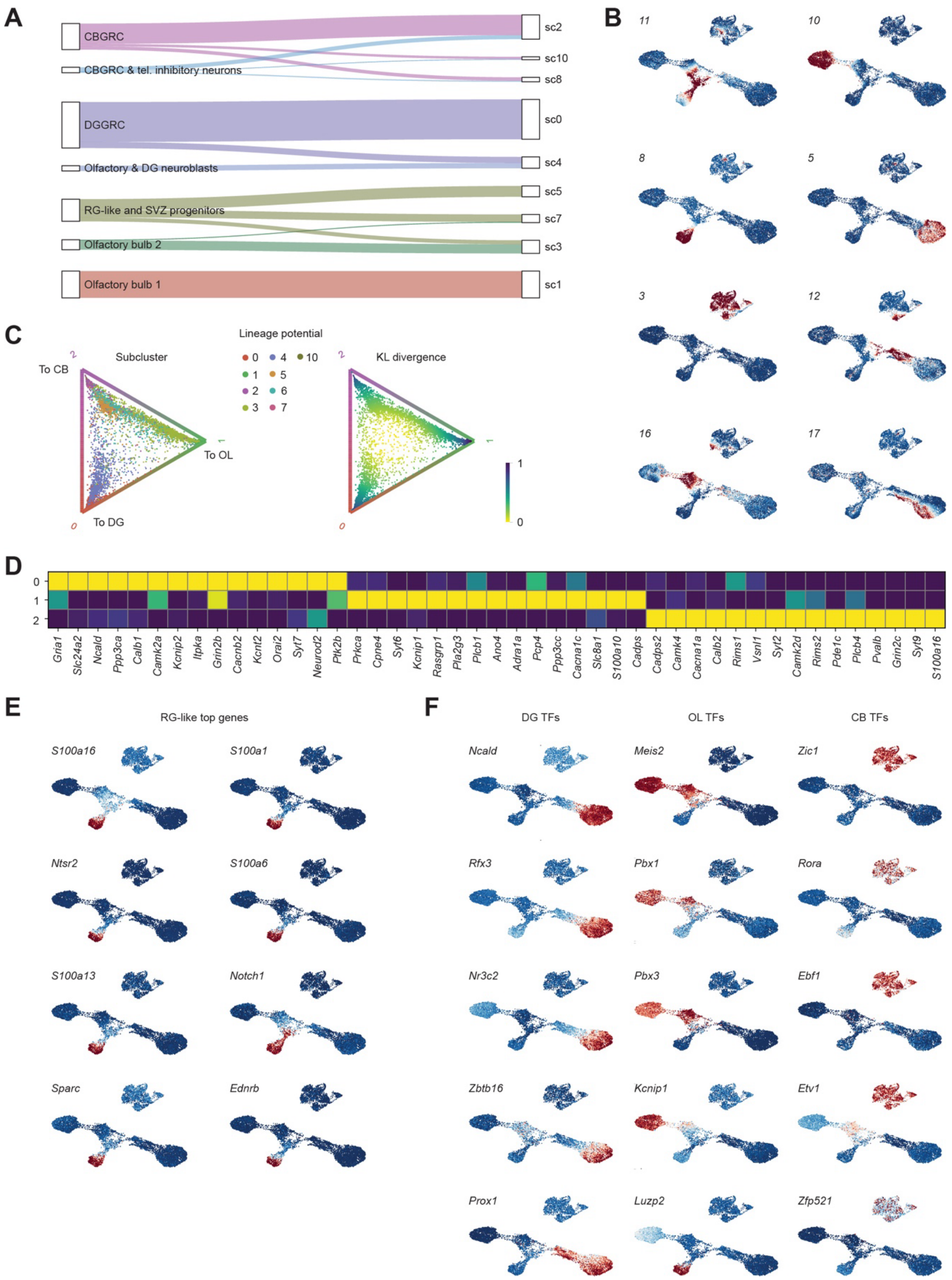

**Fig. S7. Differential gene and $Ca^{2+}$ module enrichment in postnatal developmental trajectories.** (**A**) Sankey diagram illustrative of the subclustering of the relevant CB, DG and OL $Ca^{2+}$-states. (**B**) UMAP feature plots highlighting the differential enrichment of $Ca^{2+}$ modules across the developmental subset. (**C**) Circular projection of the transition potential towards each of the three lineages colored by subcluster (left) and KL divergence (right). (**D**) Matrix plot of enriched $Ca^{2+}$ genes in each of the three terminal states. (**E**) UMAP feature plots of enriched $Ca^{2+}$ genes in the RG-like cluster. (**F**) UMAP feature plots in each of enriched TFs in each of the three lineages.

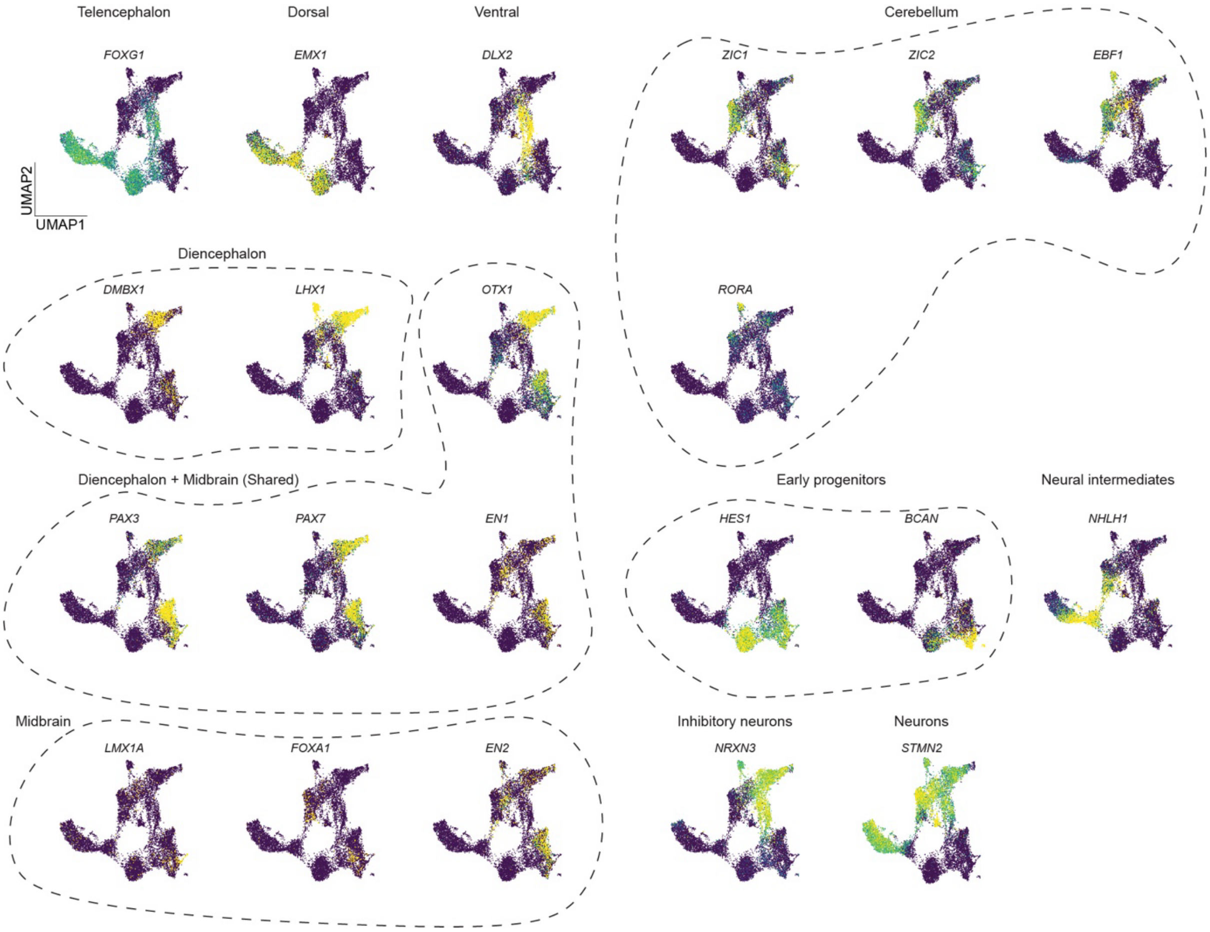

**Fig. S8. UMAP feature plots highlighting genes used to re-annotate the human developmental forebrain dataset in Van Bruggen et al. (*27*).**

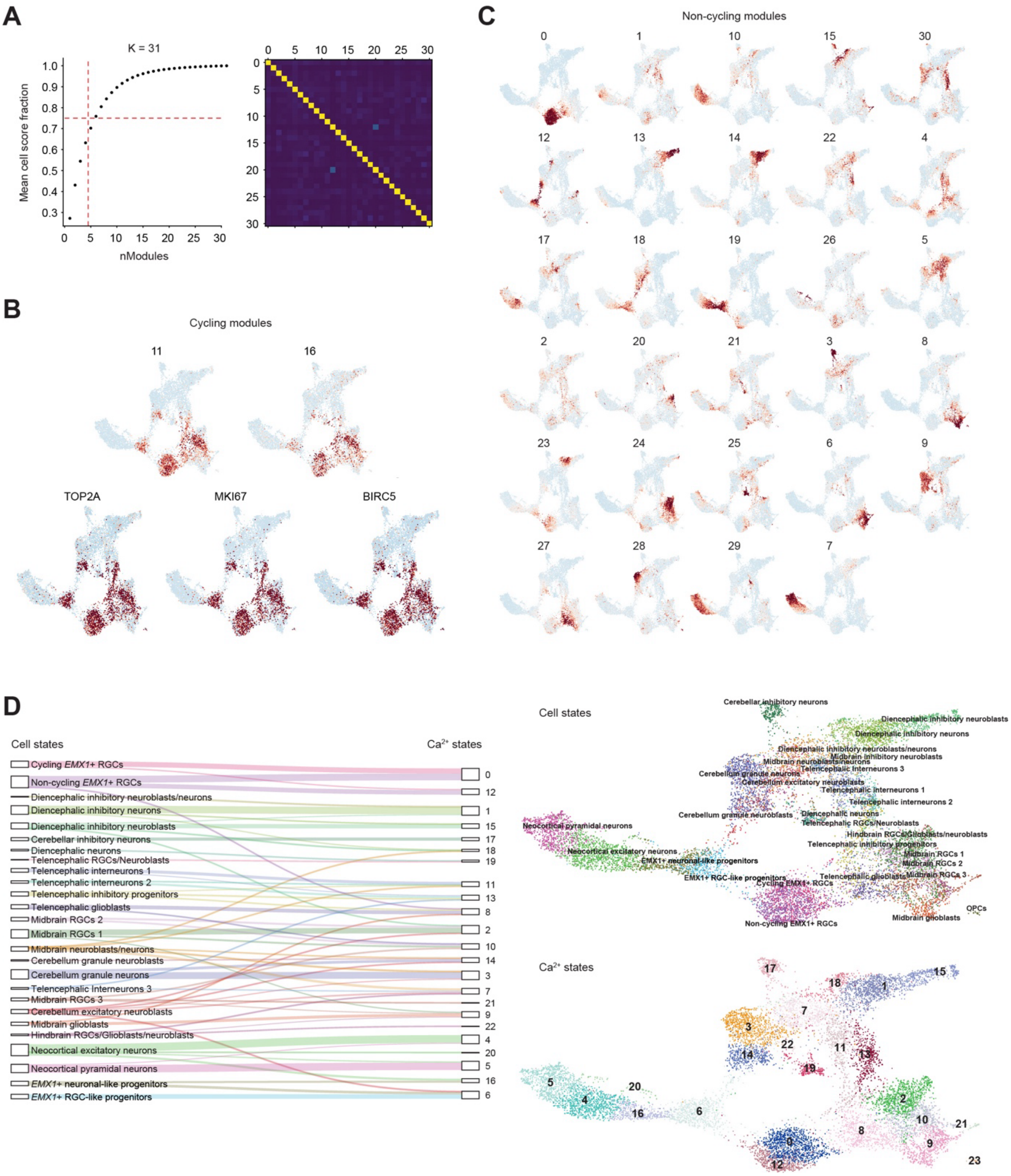

**Fig. S9. Overview of the clustering strategy used to analyze Van Bruggen et al.** (*27*)**.** (**A**) *Left*: Mean cell score fraction plotted against the number of modules. As for Zeisel et al. (*15*), scHPF was iteratively run with *K* manually tuned to the maximum value where four modules captured a mean of 70% of the cell scores. *Right:* Heatmap of the normalized mutual information scores between the modules. (**B**) UMAP plots highlighting the identified cycling modules 11 and 16 (top) and selected cycling-related genes for reference (bottom). (**C**) UMAP plots of the 29 remaining non-cycling modules, showing the differential enrichment across the dataset. (**D**) Relationship between cell and $Ca^{2+}$-states, visualized as a Sankey diagram (left) and a UMAP plot (right).

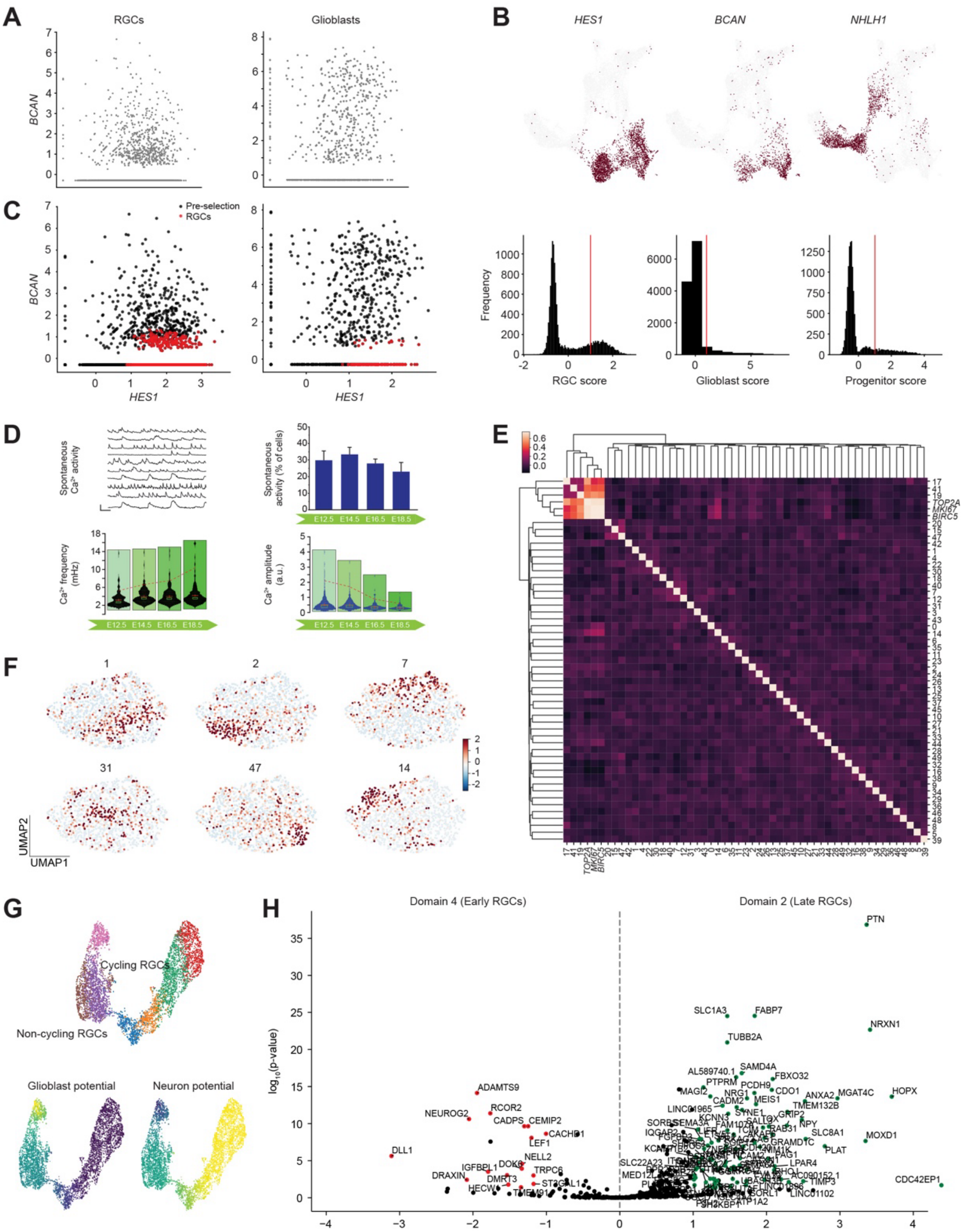

**Fig. S10. Radial glial cell heterogeneity during human development. Gene regulatory network inference using pySCENIC.** (**A**) Scatter plots of *BCAN* expression plotted against *HES1* expression in clusters defined as RGCs and glioblasts. (**B**) Binarized UMAP plots showing cells with expression of *HES1, BCAN* and *NHLH1* above gene-defined threshold cutoffs, used to conservatively select RGCs. (**C**) As in (**A**), but with selected RGCs colored in red. (**D**) Single-cell 2-photon $Ca^{2+}$ recordings from mouse cortical slices at embryonic stages E12.5, E14.5, E16.5, and E18.5 (upper left) and corresponding analyses of percent spontaneously active cells (upper right), $Ca^{2+}$ oscillatory frequency (lower left) and amplitude (lower right). (**E**) Correlation matrix of $Ca^{2+}$ modules calculated exclusively from RGCs and cycling genes. (**F**) UMAP plots showing the differential enrichment of $Ca^{2+}$ modules in the six identified functional clusters. (**G**) UMAP plots of the fate probabilities of all *EMX1+* cells towards a glioblastic and neuronal fate, respectively. (**H**) Differential expression analysis between clusters 4 (left) and 2 (right) of genes correlating with early and late radial glia.

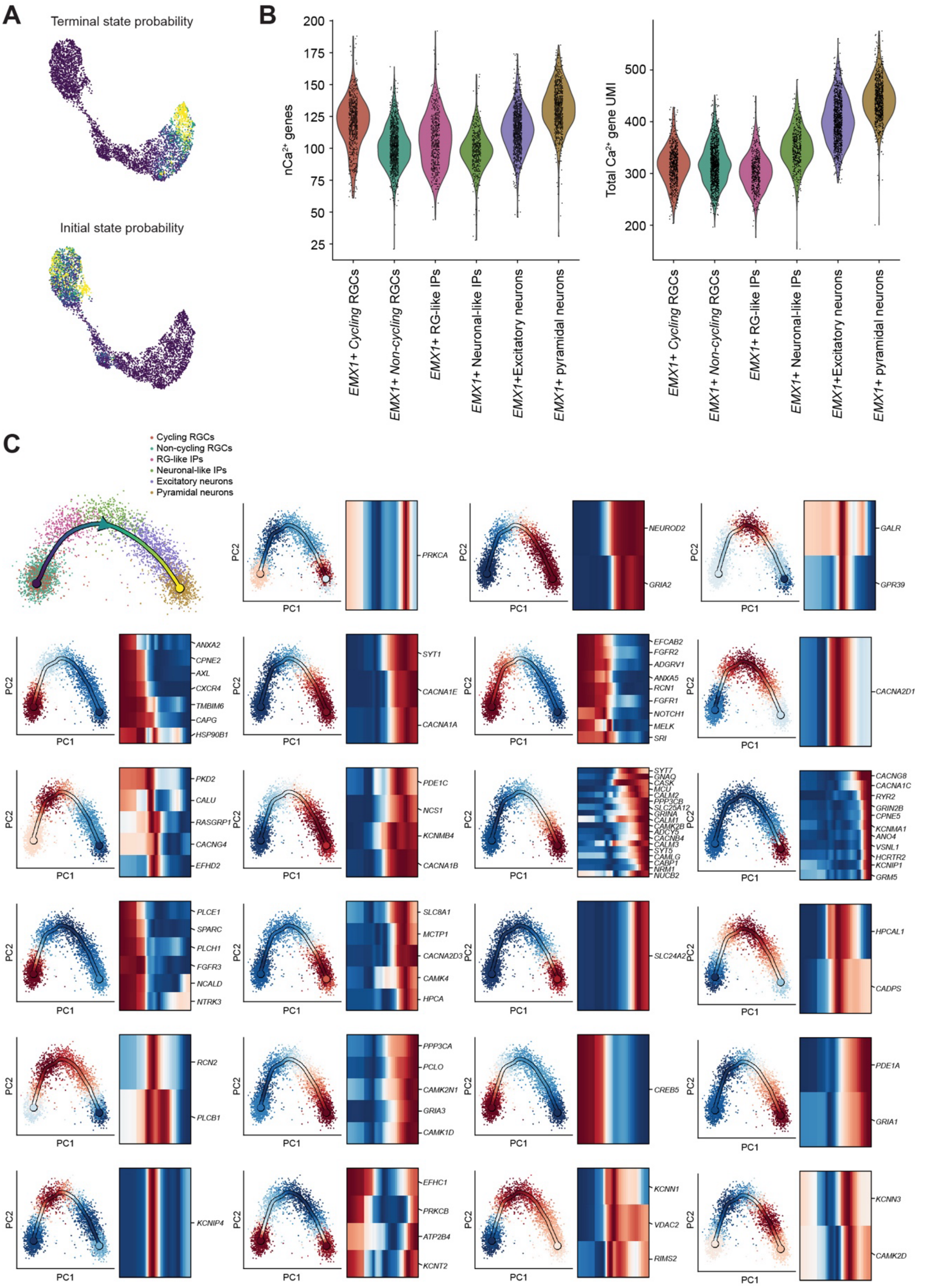

**Fig. S11. Delineation of $Ca^{2+}$ dynamics during human neocortical development.** (**A**) UMAP plots showing the initial and terminal state probabilities across the *EMX1+* lineage. (**B**) Violin plots of number of $Ca^{2+}$ genes (left) and UMI counts (right) per cell, grouped by cell states. (**C**) Scatter plots of PC1 plotted against PC2, colored by the aggregated expressions of each of the $Ca^{2+}$ gene programs, as calculated by scFates.

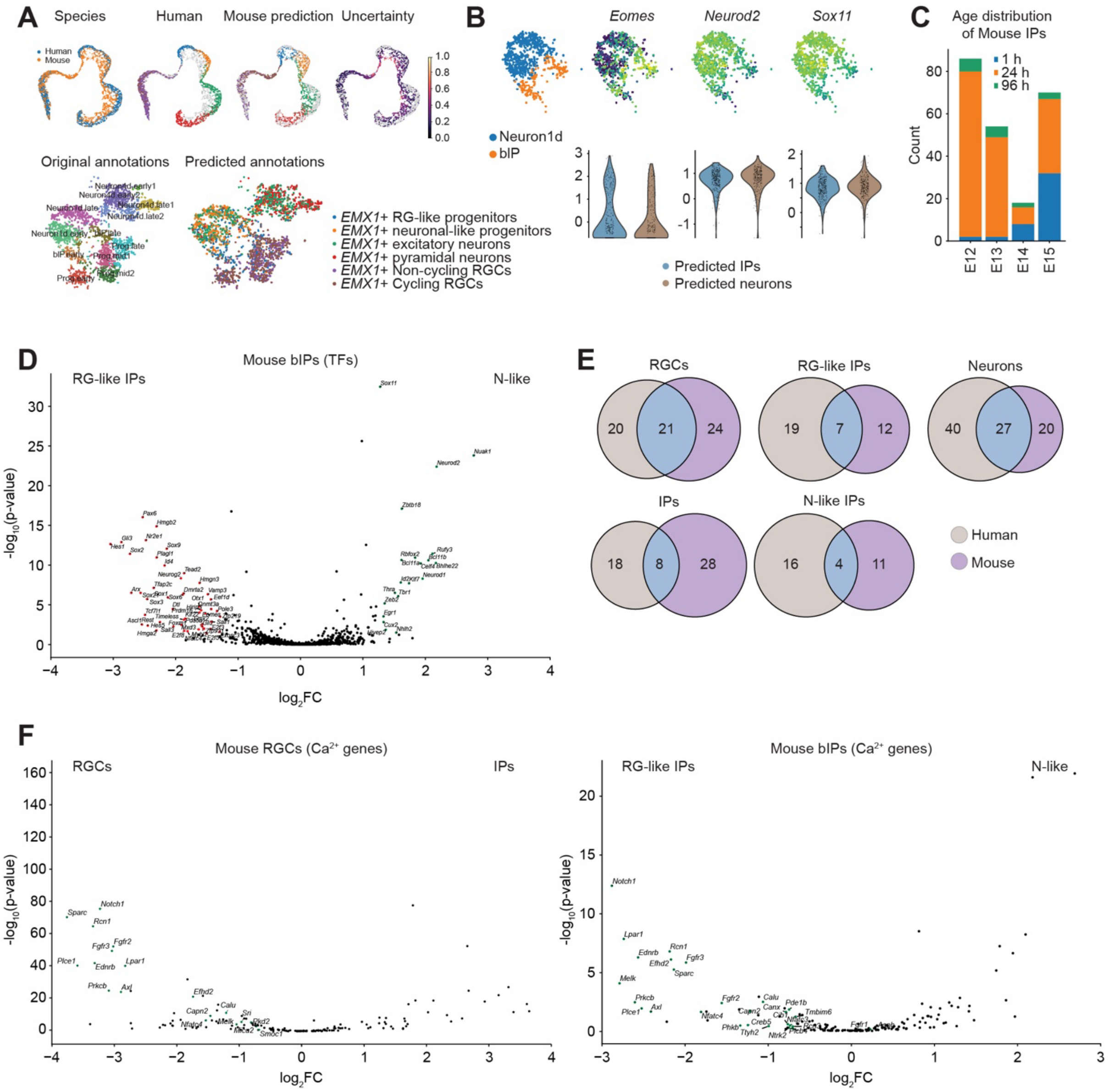

**Fig. S12. Species comparison of $Ca^{2+}$ dynamics during neocortical development.** (**A**) *Top:* Joint embedding of the human (Van Bruggen et al. (*27*)) and mouse (Telley et al. (*32*)) neocortical cells, showing the distribution of the human-defined cell states, the transferred labels in the mouse data as well as the uncertainty in the predictions. *Bottom:* TSNE highlighting the original mouse annotations and the predicted human-defined annotations. (**B**) *Top:* TSNE embedding of specifically the Neuron1d and bIP cells, as defined by the original annotations, followed by feature plots of *Eomes, Neurod2* and *Sox11*. *Bottom:* Violin plot of the same genes, grouped by the predicted human annotations. Both annotations highlight the overall similarities between the two populations. (**C**) Stacked bar plots of the age distributions of mouse IPs, grouped by embryonic age. (**D**) Volcano plot of the differentially expressed TFs in RG-like IPs (red) and neuronal-like IPs (green). (**E**) Venn diagrams showing the degree of conservation between the human-defined cell states and their predicted mouse counterparts. (**F**) *Left:* Volcano plot of the differentially expressed $Ca^{2+}$ genes in mouse RGCs (green) compared to mouse IPs. *Right:* As left, but in mouse predicted RG-like IPs versus mouse predicted N-like IPs, reflecting the similarities of the RGCs and RG-like IPs, even in mouse.
